# Predicting susceptibility to tuberculosis based on gene expression profiling in dendritic cells

**DOI:** 10.1101/104729

**Authors:** John D. Blischak, Ludovic Tailleux, Marsha Myrthil, Cécile Charlois, Emmanuel Bergot, Aurélien Dinh, Gloria Morizot, Olivia Chény, Cassandre Von Platen, Jean-Louis Herrmann, Roland Brosch, Luis B. Barreiro, Yoav Gilad

## Abstract

Tuberculosis (TB) is a deadly infectious disease, which kills millions of people every year. The causative pathogen, *Mycobacterium tuberculosis* (MTB), is estimated to have infected up to a third of the world’s population; however, only approximately 10% of infected healthy individuals progress to active TB. Despite evidence for heritability, it is not currently possible to predict who may develop TB. To explore approaches to classify susceptibility to TB, we infected with MTB dendritic cells (DCs) from putatively resistant individuals diagnosed with latent TB, and from susceptible individuals that had recovered from active TB. We measured gene expression levels in infected and non-infected cells and found hundreds of differentially expressed genes between susceptible and resistant individuals in the non-infected cells. We further found that genetic polymorphisms nearby the differentially expressed genes between susceptible and resistant individuals are more likely to be associated with TB susceptibility in published GWAS data. Lastly, we trained a classifier based on the gene expression levels in the non-infected cells, and demonstrated decent performance on our data and an independent data set. Overall, our promising results from this small study suggest that training a classifier on a larger cohort may enable us to accurately predict TB susceptibility.

## Introduction

Tuberculosis (TB) is a major public health issue. Worldwide, over a million people die of TB annually, and millions more currently live with the disease^1–3^. Successful treatment requires months of antibiotic therapy^4^, and drug-resistant strains of *Mycobacterium tuberculosis* (MTB) continuously emerge^5^. Approximately a third of the world’s population is estimated to be infected with MTB, but most are asymptomatic. While these naturally resistant individuals are able to avoid active disease, MTB might persist in a dormant state, known as latent TB^6^. In contrast, approximately 10% of individuals will develop active TB after infection with MTB^7^,^8^. Unfortunately, we are currently unable to predict if an individual is susceptible. While twin and family studies have indicated a heritable component of TB susceptibility^9–12^, genome wide association studies (GWAS) have only identified a few loci with low effect size^13–19^. Due to the highly polygenic architecture, it may be informative to examine differences between susceptible and resistant individuals at a higher level of organization, e.g. gene regulatory networks. Using this approach, previous studies have characterized gene expression profiles in innate immune cells isolated from individuals known to be susceptible or resistant to infectious diseases, including those with latent or active TB^20^ and acute rheumatic fever^21^.

We hypothesized that gene expression profiles in innate immune cells may be used to classify individuals with respect to their susceptibility to develop active TB. To test this hypothesis, we differentiated dendritic cells (DCs) from monocytes isolated from individuals that had recovered from a past episode of active TB, which we refer to as susceptible, and from individuals with confirmed latent TB, which we refer to as putatively resistant (this group is enriched in resistant individuals but we cannot exclude that some still have the potential to develop active TB^22^). We infected the DCs with MTB because these innate immune cells help shape the adaptive immune response, which is critical for fighting MTB^23,24^. We discovered that the gene expression differences between innate immune cells from resistant and susceptible individuals were present primarily in the non-infected state, that these differentially expressed genes were enriched for nearby SNPs with low p-values in TB susceptibility GWAS, and furthermore, that these gene expression levels could be used to classify individuals based on their susceptibility status.

## Results

### Susceptible individuals have an altered transcriptome in the non-infected state

We obtained whole blood samples from 25 healthy male Caucasian individuals (Supplementary Data S1). Six of the donors had recovered from active TB, and are thus putatively susceptible. The remaining 19 tested positive for latent TB without ever experiencing symptoms of active TB, and are thus putatively resistant. We isolated dendritic cells (DCs) and treated them with *Mycobacterium tuberculosis* (MTB) or a mock control for 18 hours. To measure genome-wide gene expression levels in infected and non-infected samples, we isolated and sequenced RNA using a processing pipeline designed to minimize the introduction of unwanted technical variation (Supplementary Fig. S1). We obtained a mean (± SEM) of 48 ± 6 million raw reads per sample. We performed quality control analyses to remove non-expressed genes (Supplementary Fig. S2; Supplementary Data S2), identify and remove outliers (Supplementary Fig. S3, Fig. S4, Fig. S5), and check for confounding batch effects (Supplementary Fig. S6, Fig. S7). Ultimately, data from six samples failed the quality checks and were removed from all downstream analyses (Supplementary Fig. S5).

We performed a standard differential expression analysis using a linear modeling framework (Supplementary Data S3), defined in equation (1). As expected, there was a strong response to infection with MTB in both resistant and susceptible individuals (Supplementary Fig. S8). Considering the putatively resistant individuals, we identified 3,486 differentially expressed (DE) genes between the non-infected and infected states at a q-value of 10% and an arbitrary absolute log-fold change greater than 1. Similarly, 3,789 genes were classified as DE between the non-infected and infected states in the putatively susceptible individuals. In both classes of samples, the DE genes included the important immune response factors *IL12B*, *REL*, and *TNF*. While the treatment effect was obvious in all individuals, of most interest were the patterns of gene expression differences between the putatively susceptible and resistant individuals in either the non-infected or infected states (Fig. 1). We identified 645 DE genes between putatively resistant and susceptible individuals in the non-infected state at a q-value of 10%, including *ATPV1B2*, *FEZ2*, *PSMA2*, *TNFRSF25*, and *TRIM38*. In contrast, no genes were DE between putatively resistant and susceptible individuals in the infected state (at a q-value of 10%).

**Figure 1.**
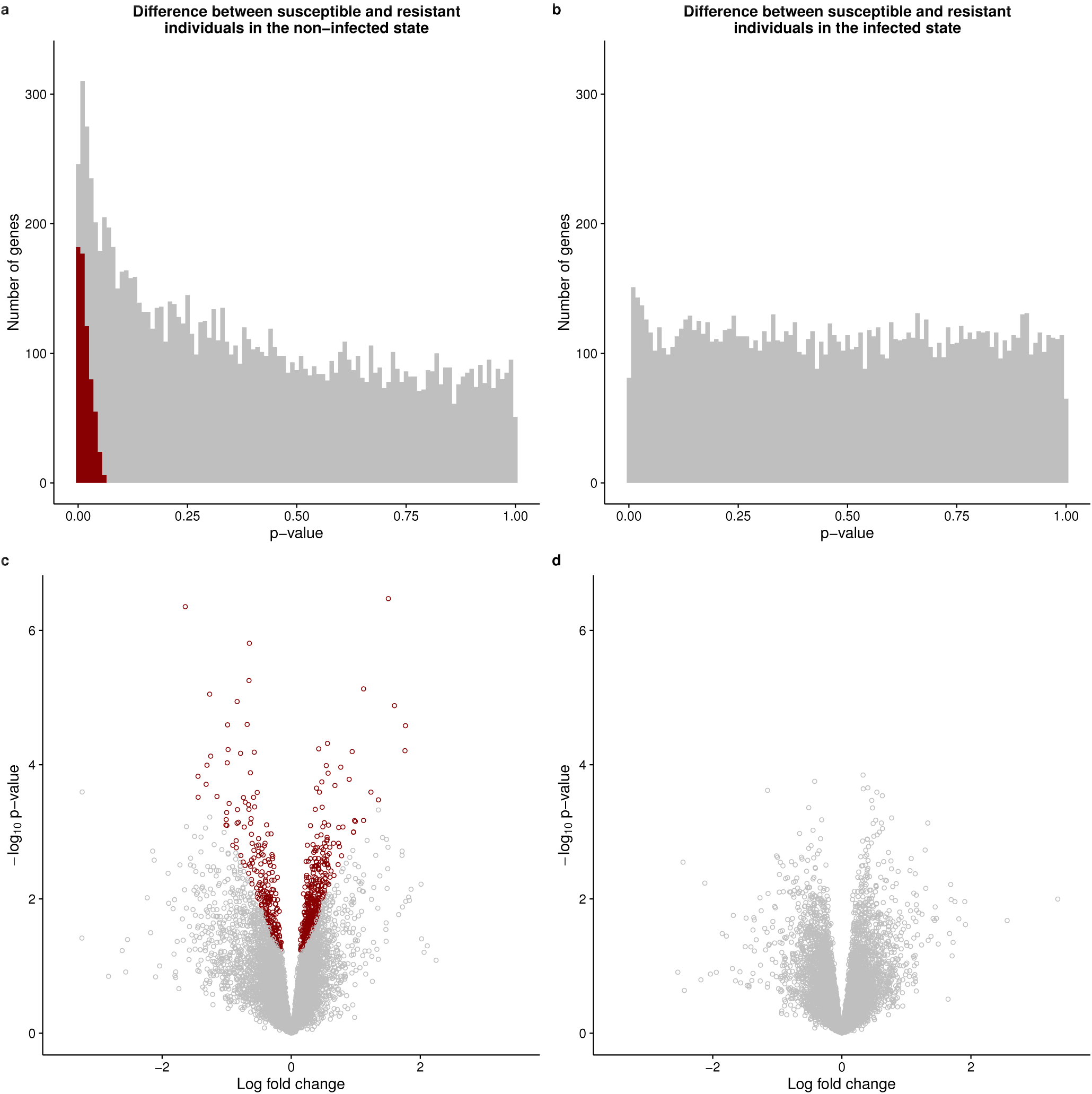
Results of differential expression analysis. The top panels show the distributions of unadjusted p-values for testing the null of no differential expression between susceptible and resistant individuals in the (a) non-infected or (b) infected state. The bottom panels show the corresponding volcano plots for the (c) non-infected and (d) infected states. The x-axis is the log fold change in gene expression level between susceptible and resistant individuals and the y-axis is the log_10_ p-value. Red indicates genes that are classified as differentially expressed with a q-value less than 10%.

### Differentially expressed genes are enriched with TB susceptibility loci

We next sought evidence that genes classified as DE in our in vitro experimental system play a role in determining susceptibility to TB. To do this, we intersected our data with results from TB susceptibility GWAS conducted in Russia^18^, The Gambia^13^, Ghana^13^, and Uganda and Tanzania^19^. We also included data from a height GWAS conducted in individuals of European ancestry^25^ as a negative control. To perform a combined analysis of our gene expression data and the GWAS results, we had to define pairs of genes (for which we have expression data) and SNPs (for which we obtained GWAS *P* values). Thus, each gene in our expression data was coupled with the GWAS SNP with the lowest p-value among all tested SNPs located within 50 kb of the gene’s transcription start site (Supplementary Data S4; this gene-SNP definition was performed separately for each GWAS data set).

Once we defined gene–SNP pairs, we asked whether differences in gene expression levels between putatively resistant and susceptible individuals could help us identify genetic variation that is associated with susceptibility to TB. In other words, we asked whether increasing evidence for DE genes is associated with low GWAS p-values. To do so, we calculated the fraction of SNPs with a GWAS p-value lower than 0.05 among SNPs that were paired with ranked subsets of genes whose expression profiles show increasing effect size of expression differences between putatively resistant and susceptible individuals. In order to assess the significance of the observations, we performed 100 permutations of the enrichment analysis to derive an empirical p-value.

Using this approach, we observed a clear enrichment (empirical P < 0.01) of low p-values for TB GWAS SNPs that are paired with genes that are differentially expressed between susceptible and resistant individuals in the non-infected state (Fig. 2a). In fact, we observed significant enrichments of lower GWAS p values (empirical P < 0.01) in all 4 TB susceptibility GWAS (Russia, The Gambia, Ghana, Uganda and Tanzania) (Supplementary Fig. S10) for all 4 differential expression contrasts, namely resistant vs. susceptible individuals in the non-infected state (Fig. 2c), resistant vs. susceptible individuals in the infected state (Fig. 2d), effect of treatment in resistant individuals (Fig. 2e), and effect of treatment in susceptible individuals (Fig. 2f). Reassuringly, we did not observe an enrichment of low p values (empirical P > 0.01) when we used the same approach to consider data from the height GWAS (Fig. 2bcdef).

**Figure 2.**
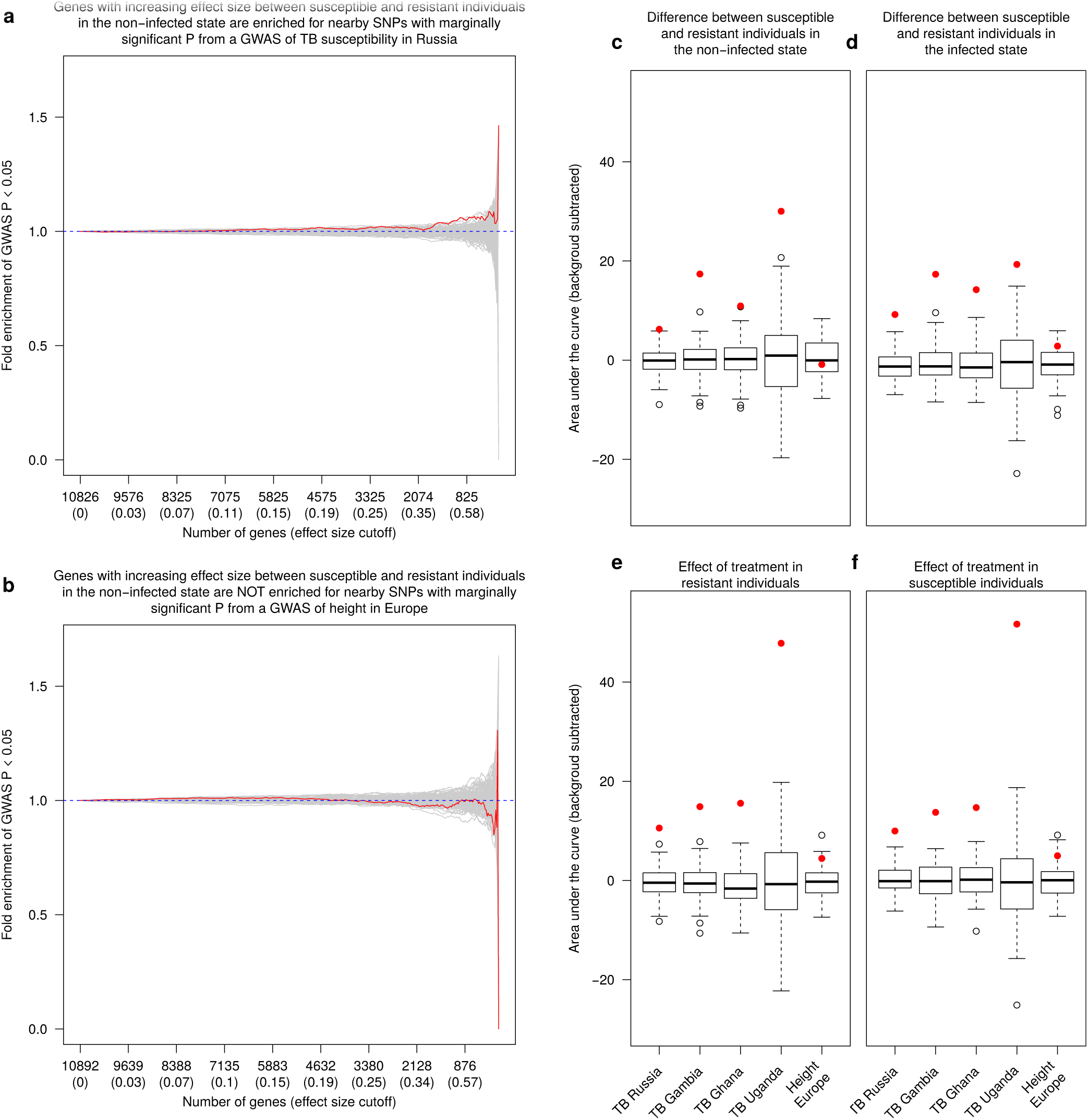
Comparison of differential expression and TB susceptibility GWAS results. (a and b) The y-axis is the fold enrichment of SNPs with p-value less than 0.05 from the (a) GWAS of TB susceptibility in Russia^18^ or (b) height in individuals of European descent^25^. The x-axis is bins of genes with increasingly stringent effect size cutoffs of the absolute expression log fold change between putatively susceptible and resistant individuals in the non-infected state. The effect size cutoffs were chosen such that each bin from left to right contained approximately 25 fewer genes. The red line shows the results from the actual data. The grey lines are the results from 100 permutations. The dashed blue line at y=1 represents the null expectation. (c-f) Boxplots of the area under the curve of the fold enrichment (red line in a and b) minus the background level (blue y = 1 line in a and b) for each of the 5 GWAS^13^, ^18^, ^19^ considered for the 4 differential expression contrasts: (c) resistant vs. susceptible individuals in the non-infected state, (d) resistant vs. susceptible individuals in the infected state, (e) effect of treatment in resistant individuals, (f) effect of treatment in susceptible individuals. The boxplot is the result of the 100 permutations, and the red point is the result from the actual data. As a reference, the leftmost boxplot in (c) corresponds to the enrichment plot in (a), and the rightmost boxplot in (c) corresponds to the enrichment plot in (b).

### Susceptibility status can be predicted based on gene expression data

Next we attempted to build a gene expression-based classifier to predict TB susceptibility status (Supplementary Data S5). We focused on the gene expression levels measured in the non-infected state both because this is where we observed the largest gene regulatory differences between putatively susceptible and resistant individuals (Fig. 1ac), and also because, from the perspective of an ultimate translational application, it is more practical to obtain gene expression data from non-infected DCs. We trained a support vector machine using the 99 genes that were differentially expressed between resistant and susceptible individuals in the non-infected state at a q-value less than 5% (see Methods for a full description of how we selected this model). Encouragingly, we observed a clear separation between putatively susceptible and resistant individuals when comparing the predicted probability of being susceptible to TB for each sample obtained from leave-one-out-cross-validation (Fig. 3a). Using a cutoff of 0.25 for the predicted probability of being susceptible to TB, we obtained a sensitivity of 100% (5 out of 5 susceptible individuals classified as susceptible), a specificity of ~88% (15 out of 17 resistant individuals classified as resistant), and a positive predictive value (PPV) of ~71% (5 of 7 individuals classified as susceptible were susceptible).

**Figure 3.**
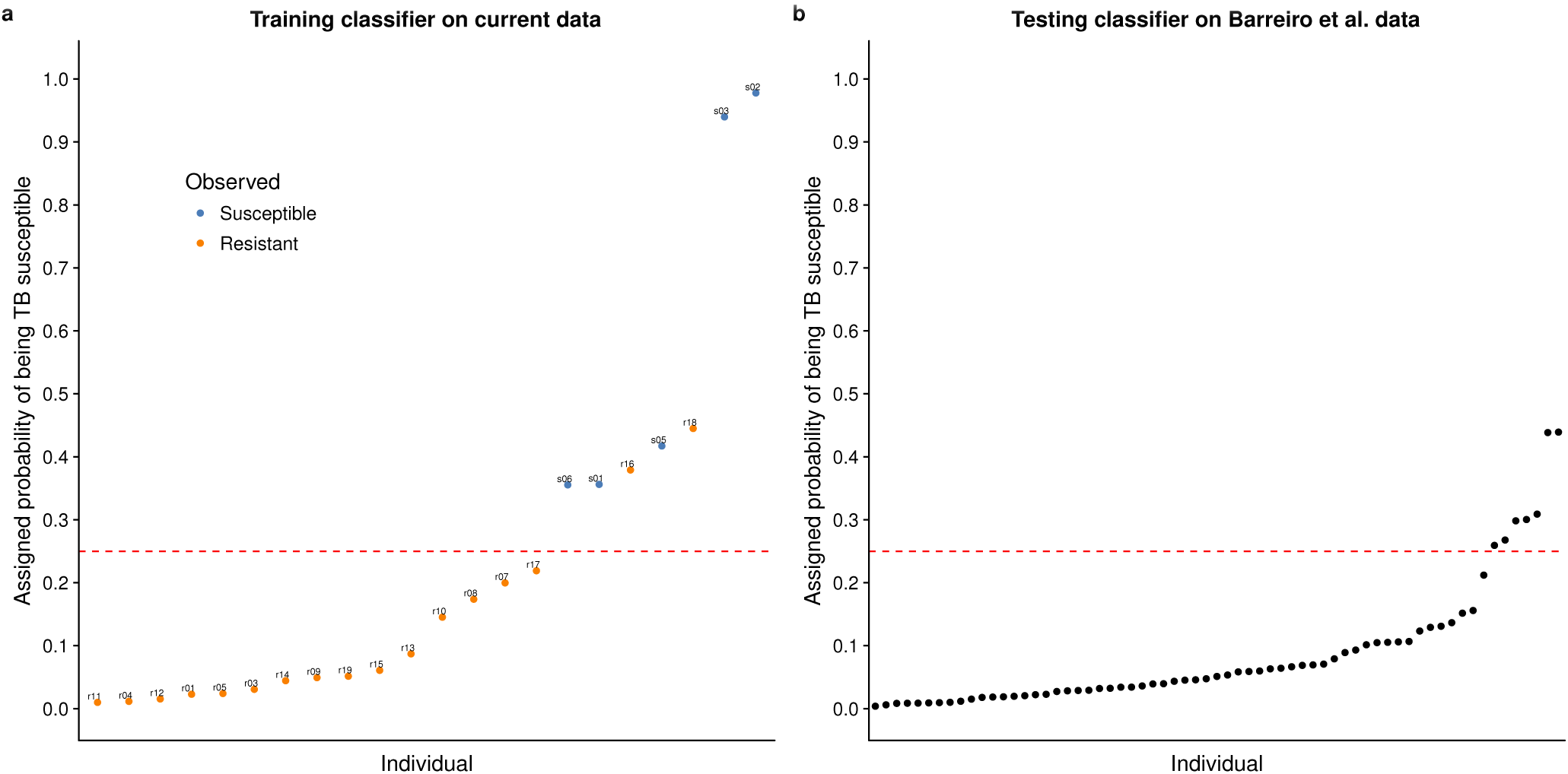
Classifying TB susceptible individuals using a support vector machine model. (a) The estimates of predicted probability of TB susceptibility from the leave-one-out-cross-validation for individuals in the current study. The blue circles represent individuals known to be susceptible to TB, and orange those resistant to TB. The horizontal dashed red line at a probability of 0.25 separates susceptible and resistant individuals. (b) The estimates of predicted probability of TB susceptibility from applying the classifier trained on the data from the current study to a test set of independently collected healthy individuals^24^.

Unfortunately our current data set is too small to properly split into separate training and testing sets (it is challenging to collect samples from previous TB patients, who are healthy and have no medical reason to go back for a GP visit). To our knowledge, there are also no other suitable data sets available with which to test out classifier (that said, see Supplementary Fig. S17 for the results of applying the classifier to a non-ideal data set, which measured gene expression in macrophages from a small number of individuals^20^). Thus, in order to further assess the plausibility of our model, we applied the classifier to data from an independent study, which collected genome-wide gene expression levels in DCs from 65 healthy individuals^24^, none with a previous history of TB. Using the cutoff of 0.25 for the probability of being susceptible to TB (determined to be optimal in the training set), ~11% (7 of 65) of the individuals were classified as susceptible to TB (Fig. 3b). Adjusting for the PPV obtained from the training set (~71%), our model predicted that ~7.7% of the healthy individuals were susceptible. While we cannot confirm this result (the true susceptibility status of these 65 individuals is unknown), this observation is encouraging because our estimate is similar to the commonly used inference that roughly 10% of the general population is susceptible to TB.

## Discussion

We obtained dendritic cells (DCs) from individuals that were known to be putatively susceptible or resistant to developing active tuberculosis (TB) and measured genome-wide gene expression levels in non-infected DCs and DCs infected with *Mycobacterium tuberculosis* (MTB) for 18 hours. As expected, there were large changes in gene expression due to MTB infection in the DCs from both putatively resistant and susceptible individuals (Supplementary Fig. S8). We identified 645 genes which were differentially expressed (DE) between susceptible and resistant individuals in the non-infected state; whereas, we did not observe any DE genes between susceptible and resistant individuals in the infected state (Fig. 1). This suggests that the differences in the transcriptomes between DCs of resistant and susceptible individuals are present pre-infection. Yet, 18 hours after infection gene expression profiles in both susceptible and resistant individuals have converged to a similar gene regulatory network, presumably to fight the infection. We confirmed that the absence of DE genes in the infected state is **not** caused by a decrease in statistical power due to an overall increase in gene expression variance upon infection (Supplementary Fig. S9). We chose to measure gene expression 18 hours post-infection because this time point was previously associated with a large change in genome-wide gene expression levels^26^. Given our observations, however, future studies investigating the difference in the innate immune response between individuals resistant and susceptible to TB may want to focus on earlier time points post-infection.

It is important to note that our study was not designed to uncover the mechanisms underlying susceptibility or resistance to TB, but to try and find a gene regulatory signature that might allow us to classify individuals as either susceptible or resistant. That said, among the 645 DE genes between resistant and susceptible individuals in the non-infected state, there were many interesting genes involved in important innate immune activities critical for fighting MTB and other pathogens such as autophagy^27,28^, phagolysosomal acidification, and antigen processing. In particular, *FEZ2*, a suppressor of autophagosome formation^29^, was down-regulated when DCs were infected with MTB; however, in the non-infected DCs, this gene has elevated expression level in susceptible compared with resistant individuals. In turn, *ATP6V1B2*, a gene coding for a subunit of the proton transporter responsible for acidifying phagolysosomes^30–32^, has increased expression in susceptible individuals compared to resistant in the non-infected state. Lastly, genes coding for nine subunits of the proteasome, which is critical for processing of MTB antigens to be presented via major histocompatibility complex (MHC) class I molecules^33–36^, have increased expression in susceptible individuals compared to resistant in the non-infected state. These genes are candidates for future functional studies investigating the mechanisms of TB susceptibility.

We observed that DE genes in our in *vitro* experimental system were enriched for lower GWAS p-values (Fig. 2). This suggests that such *in vitro* approaches are informative for interrogating the genetic basis of disease susceptibility. That being said, we recognize a major caveat with this analysis is that assigning SNPs to their nearest gene on the linear chromosome is problematic because regulatory variants can have longer range effects. Nevertheless, considering this limitation, it was encouraging that we were able to detect evidence of the genetic basis of TB susceptibility in this system.

Not only did this analysis identify a global enrichment of TB susceptibility loci, but by intersecting the expression and GWAS data, we were able to identify a few interesting candidate genes, which were only marginally significant in the original GWAS (Supplementary Data S4). Here we highlight two genes (*CCL1* and *UNC13A*), which have been previously shown to play important roles in MTB infection. *CCL1* is a chemokine that stimulates migration of monocytes^37^. In our study, it was upregulated in susceptible individuals compared to resistant in both the non-infected and infected states (but did not reach statistical significance in either) and was statistically significantly upregulated with MTB treatment. Furthermore, the nearby SNP assigned to *CCL1* had a p-value less than 0.01 in the TB susceptibility GWAS from The Gambia and Ghana. A previous differential expression study of TB susceptibility (discussed in more detail below) found that *CCL1* was upregulated to a greater extent 4 hours post-infection with MTB in macrophages isolated from individuals with active TB (i.e. susceptible) compared to individuals with latent TB (i.e. resistant)^20^. Additionally they performed a candidate gene association study and found that SNPs nearby *CCL1* were associated with TB susceptibility. In our previous study, we discovered that *CCL1* was one of only 288 genes that were differentially expressed in macrophages 48 hours post-infection with MTB and related mycobacterial species but not unrelated virulent bacteria^38^. *UNC13A* is involved in vesicle formation^39^. In our study, it was downregulated in susceptible individuals compared to resistant in both the non-infected and infected states (but did not reach statistical significance in either) and was statistically significantly upregulated with MTB treatment. Furthermore, the nearby SNP assigned to *UNC13A* had a p-value less than 0.01 in the TB susceptibility GWAS from Russia, The Gambia, and Ghana. In our past study mapping expression quantitative trait loci (eQTLs) in DCs 18 hours post-infection with MTB, *UNC13A* was one of only 98 genes which were associated with an eQTL post-infection but not pre-infection, which we called MTB-specific eQTLs^24^. Thus our new results increased the evidence that *CCL1* and *UNC13A* play important roles in TB susceptibility.

Previous attempts to use gene expression based classifiers in the context of TB have focused on predicting the status of an infection rather than the susceptibility status of an individual^8,40,41^. In other words, the goal of most previous studies was to detect individuals in the early stages of active TB when antibiotic intervention would be most effective or to monitor the effectiveness of a treatment regimen^42^. In contrast, our goal was not to distinguish between active and latent TB, but instead to be able to determine susceptibility status before individuals are infected with MTB. Even with our small sample size, we were able to successfully train a classier with high sensitivity and decent specificity. Because such a classification of susceptibility status could affect the decision of whether or not to take antibiotics to treat latent TB^6^, false negatives (susceptible individuals mistakenly classified as resistant) would be much more harmful than false positives (resistant individuals mistakenly classified as susceptible). For that reason, we emphasized sensitivity over specificity.

To our knowledge, our study was only the second to collect data from *in vitro* MTB-infected innate immune cells isolated from individuals known to be putatively susceptible to MTB (Thuong et al., 2008). However, there were substantial differences between our study and that of Thuong et al., 2008^20^. First, they derived and infected macrophages, the primary target host cell in which MTB resides; whereas, we derived and infected DCs, which play a larger role in stimulating the adaptive immune response to MTB. Second, we collected samples from a larger number of putatively resistant individuals (19 versus 4), increasing our power to distinguish between the gene expression profiles of susceptible and resistant individuals. Third, they measured gene expression with microarrays; whereas, we used RNA-sequencing. Considering the substantial technical differences between the methods used and the biological differences between DCs and macrophages^26,43^, unsurprisingly, we were unable to identify the susceptible individuals from Thuong et al., 2008^20^ using our classifier (Supplementary Fig. S17).

Indeed, at this time, we are not aware of any other data set from healthy individuals known to be sensitive to TB, with which we can further test our classifier. When we applied our classifier to an independent set of non-infected DCs isolated from healthy individuals of unknown susceptibility status, our model predicted that ~7.7-11% of the individuals were susceptible to TB, which reassuringly is similar to the average in the general population (10%). Despite this, our results must be interpreted cautiously; at best as a proof-of-principle, due to our very small sample size of only 5 susceptible individuals. That said, our promising results in this small study suggest that collecting blood samples from a larger cohort of susceptible individuals would enable building a gene expression based classifier able to confidently assess risk of TB susceptibility. By reducing the number of resistant individuals receiving treatment for latent TB, we can eliminate the adverse health effects of a 6 month regimen of antibiotics for these individuals and also reduce the selective pressures on MTB to develop drug resistance.

## Methods

### Ethics statement

We recruited 25 subjects to donate a blood sample for use in our study. All methods were carried out in accordance with relevant guidelines and regulations. All participants gave written informed consent in accordance with the Declaration of Helsinki principles. Peripheral human blood was collected from patients at ICAReB platform of Institut Pasteur Paris and at the Centre for Infectious Disease Prevention, University hospital Caen. The Protocol has been approved by French Ethical Committee (CPP North Ouest III, n° A12 - D33-VOL.13), and by the Institutional Review Boards of the University of Chicago (10-504-B) and the Institut Pasteur (IRB00006966).

### Sample collection

We collected whole blood samples from healthy Caucasian male individuals living in France. The putatively resistant individuals tested positive for latent TB in an interferon-*γ* release assay, but had never developed active TB. The putatively sensitive individuals had developed active TB in the past, but were currently healthy.

### Isolation and infection of dendritic cells

We performed these experiments as previously described^24^. Briefly, we isolated mononuclear cells from the whole blood samples using Ficoll-Paque centrifugation, extracted monocytes via CD14 positive selection, and differentiated the monocytes into dendritic cells (DCs) by culturing them for 5 days in RPMI 1640 (Invitrogen) supplemented with 10% heat-inactivated FCS (Dutscher), L-glutamine (Invitrogen), GM-CSF (20 ng/mL; Immunotools), and IL-4 (20 ng/mL; Immunotools). Next we infected the DCs with *Mycobacterium tuberculosis* (MTB) H37Rv at a multiplicity of infection of 1-to-1 for 18 hours.

### RNA extraction and sequencing

We extracted RNA using the Qiagen miRNeasy Kit and prepared sequencing libraries using the Illumina TruSeq Kit. We sent the master mixes to the University of Chicago Functional Genomics Facility to be sequenced on an Illumina HiSeq 4000. We designed the batches for RNA extraction, library preparation, and sequencing to balance the experimental factors of interest and thus avoid potential technical confounders (Supplementary Fig. S1).

### Read mapping

We mapped reads to human genome hg38 (GRCh38) using Subread^44^ and discarded non-uniquely mapping reads. We downloaded the exon coordinates of 19,800 Ensembl^45^ protein-coding genes (Ensembl 83, Dec 2015, GRCh38.p5) using the R/Bioconductor^46^ package biomaRt^47,48^ and assigned mapped reads to these genes using featureCounts^49^.

### Quality control

First we filtered genes based on their expression level by removing all genes with a transformed median log_2_ counts per million (cpm) of less than zero. This step resulted in a set of 11,336 genes for downstream analysis (Supplementary Fig. S2, Supplementary Data S2). Next we used principal components analysis (PCA) and hierarchical clustering to identify and remove 6 outlier samples (Supplementary Fig. S3, Fig. S4, Fig. S5). We did this systematically, by removing any sample whose data projections did not fall within two standard deviations of the mean for any of the first six PCs (for the first PC, which separated the samples by treatment, we calculated a separate mean for the non-infected and infected samples).

After filtering lowly expressed genes and removing outliers, we performed the PCA again to check for any potential confounding technical batch effects (Supplementary Fig. S6). Reassuringly, the major sources of variation in the data were from the biological factors of interest. PC1 was strongly correlated with the effect of treatment, and PCs 2-6 were correlated with inter-individual variation. The only concerning technical factor was the infection experiments, which were done in 12 separate batches (Supplementary Fig. S1). Infection batch correlated with PCs 3 and 5; however, we verified that this variation was not confounded with our primary outcome of interest, TB susceptibility (Supplementary Fig. S7).

### Differential expression analysis

We used limma+voom^50–52^ to implement the following linear model to test for differential expression:

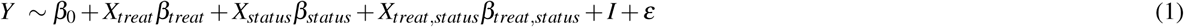

Where *β*_0_ is the mean expression level in non-infected cells of resistant individuals, *β_treat_* is the fixed effect of treatment in resistant individuals, *β_status_* is the fixed effect of susceptibility status in non-infected cells, *β_treat;status_* is the fixed interaction effect of treatment in susceptible individuals (i.e. modeling the interaction between treatment and susceptibility status), and I is the random effect of individual. The random individual effect was implemented using the limma function duplicateCorrelation^53^. To jointly model the data with voom and duplicateCorrelation, we followed the recommended best practice of running both voom and duplicateCorrelation twice in succession^54^.

We used the model to test different hypotheses (Supplementary Data S3). We identified genes which were differentially expressed (DE) between infected and non-infected DCs of resistant individuals by testing *β_treat_* = 0, genes which were DE between infected and non-infected DCs of susceptible individuals by testing *β_treat_* + *β_treat;status_* = 0, genes which were DE between susceptible and resistant individuals in the non-infected state by testing *β_status_* = 0, and genes which were DE between susceptible and resistant individuals in the infected state by testing *β_status_* + *β_treat;status_* = 0. We corrected for multiple testing using q-values estimated via adaptive shrinkage^55^ and considered differentially expressed genes as those with a q-value less than 10%.

Note that we also tested the interaction term, *β_treat;status_* = 0, to identify genes in which the difference in expression level between the infected and non-infected states was significantly different between susceptible and resistant individuals. However, as expected since no DE genes were identified between susceptible and resistant individuals in the infected state (see Results), the results of testing the interaction term were partially redundant with the results of testing differences between susceptible and resistant individuals in the non-infected state, and thus we ignored these results throughout this study.

### Combined analysis of gene expression data and GWAS results

The GWAS p-values were from previously published studies of TB susceptibility conducted in Russia^18^, The Gambia^13^, Ghana^13^, and Uganda and Tanzania^19^ (and a height GWAS in individuals of European descent^25^). To perform a combined analysis of the gene expression and the summary statistics from each GWAS, we assigned each gene to the SNP with the minimum GWAS p-value out of all the SNPs located within 50 kb up or downstream of its transcription start site. Specifically, we obtained the genomic coordinates of the SNPs with the R/Bioconductor^46^ package SNPlocs.Hsapiens.dbSNP144.GRCh38 and matched SNPs to nearby genes using GenomicRanges^56^. 10,265 to 11,060 of the 11,336 genes were assigned an association p-value depending on the GWAS (Supplementary Data S4). For each of the 4 differential expression contrasts we tested (resistant vs. susceptible individuals in the non-infected state, resistant vs. susceptible individuals in the infected state, effect of treatment in resistant individuals, effect of treatment in susceptible individuals), we also performed an enrichment analysis. To do so, we calculated the fraction of genes assigned a GWAS SNP with p-value less than 0.05 for bins of genes filtered by increasingly stringent cutoffs for the observed differential expression effect size (the absolute value of the log fold change). The effect size cutoffs were chosen such that on average each subsequent bin differed by 25 genes. To measure enrichment, we calculated the area under the curve using the R package flux^57^ (we also subtracted the background area under the line y = 1 because the number of genes assigned a SNP varied across the GWAS). In order to assess significance, we calculated the area under the curve for 100 permutations of the data. All differential expression tests were statistically significantly enriched for SNPs with low GWAS p-values in every TB susceptibility GWAS (empirical *P* < 0.01) and not enriched for the height GWAS (empirical *P* > 0.01) (Fig. 2; Supplementary Fig. S10).

### Classifier

The training set included data from the 44 high-quality non-infected samples from this study with known susceptibility status. The test set included the 65 non-infected samples from one of our previous studies in which the susceptibility status is unknown^24^, and thus assumed to be similar to that in the general population (~10%) (we also tested the classifier on data from a small study of macrophages^20^, Supplementary Fig. S17). Because the two studies are substantially different, we took multiple steps to make them comparable. First, we subset to include only those 9,450 genes which were assayed in both. Second, because the dynamic range obtained from RNA-seq (current study) and microarrays (previous study^24^) were different, we normalized the gene expression levels to a standard normal (*µ* = 0, σ = 1) distribution (Supplementary Fig. S11; note however that this strategy is unable to correct for the inability of microarrays to accurately quantify genes with expression levels that result in fluorescence levels below the background level or above the saturation limit). Third, we corrected for the large, expected batch effect between the two studies by regressing out the first PC of the combined expression data using the limma function removeBatchEffect^52^ (Supplementary Fig. S12).

To identify genes to use in the classifier, we performed a differential expression analysis on the normalized, batch-corrected data from the current study using the same approach described above (with the exception that we no longer used voom^51^ since the data were no longer counts). Specifically, we tested for differential expression between susceptible and resistant individuals in the non-infected state and identified sets of genes to use in the classifier by varying the q-value cutoff. Cutoffs of 5%, 10%, 15%, 20%, and 25% corresponded to gene set sizes of 99, 385, 947, 1,934, and 3,697, respectively. We used the R package caret^58^ to train 3 different machine learning models: elastic net^59^, support vector machine^60^, and random forest^61^ (the parameters for each individual model were selected using the Kappa statistic). To assess the results of the model on the training data, we performed leave-one-out-cross-validation (LOOCV). In order to choose the model with the best performance, we calculated the difference between the mean of the LOOCV-estimated probabilities of being TB resistant for the samples known to be TB resistant and the corresponding mean for the samples known to be TB susceptible. This metric emphasized the ability to separate the susceptible and resistant individuals into two separate groups. Using this metric, the best performing model was the support vector machine with the 99 genes that are significantly differentially expressed at a q-value of 5% (Fig.3a, Supplementary Fig. S13, Supplementary Data S5); however, both the elastic net (Supplementary Fig. S14) and random forest (Supplementary Fig. S15) had similar performance. Lastly, we tested the classifier by predicting the probability of being TB susceptible in the 65 healthy samples (Fig. 3b). For evaluating the predictions on the test set of individuals with unknown susceptibility status, we used a relaxed cutoff of the probability of being TB susceptible of 0.25, which was based on the ability of the model at this cutoff to classify all TB susceptible individuals in the training set as susceptible with only 2 false positives. As expected, the 99 genes used in the classifier had similar normalized, batch-corrected median expression levels in the non-infected state across both studies (Supplementary Fig. S16).

### Software implementation

We automated our analysis using Python (https://www.python.org/) and Snakemake^62^. Our processing pipeline used the general bioinformatics software FastQC (http://www.bioinformatics.babraham.ac.uk/projects/fastqc/), MultiQC^63^, samtools^64^, and bioawk (https://github.com/lh3/bioawk). We used R^65^ for all statistics and data visualization. We obtained gene annotation information from the Ensembl^45^ and Lynx^66^ databases. The computational resources were provided by the University of Chicago Research Computing Center. All code is available for viewing and reuse at https://github.com/jdblischak/tb-suscept.

### Data availability

The raw fastq files have been deposited in NCBI’s Gene Expression Omnibus^67^ and are accessible through GEO Series accession number GSE94116 (http://www.ncbi.nlm.nih.gov/geo/query/acc.cgi?acc=GSE94116). The RNA-seq gene counts and other summary data sets are included as Supplementary Data and are also available for download at https://github.com/jdblischak/tb-suscept.

While a small subset of the GWAS summary statistics required for partially reproducing our results were included in Supplementary Data S4, we do not have permission to share the full set of summary statistics from the previously published TB susceptibility GWAS that would be required for fully reproducing our results. To access the summary statistics, contact the authors directly: Russia - Sergey Nejentsev, Ghana and The Gambia - Thorsten Thye, Uganda and Tanzania - Scott M. Williams. The summary statistics for the height GWAS can be downloaded from the GIANT Consortium’s website (http://portals.broadinstitute.org/collaboration/giant/index.php/GIANT_consortium_data_files).

## Acknowledgements

We thank Matthew Stephens and John Novembre for providing feedback and Gilad lab members for helpful discussion. We thank Marie-Noëlle Ungeheuer and Nathalie Jolly for help recruiting subjects. We thank Sergey Nejentsev for sharing data from the GWAS in Russia. We thank Thorsten Thye for sharing data from the GWAS in Ghana and The Gambia. We thank Rafal S. Sobota, Catherine M. Stein, Giorgio Sirugo, and Scott M. Williams for sharing data from the GWAS in Uganda and Tanzania. This study was funded by National Institutes of Health (NIH) Grant AI087658 to Y.G. and L.T.J.D.B. was supported by NIH Grant T32GM007197. The content is solely the responsibility of the authors and does not necessarily represent the official views of the NIH.

## Author contributions

Y.G., L.T., and L.B.B. conceived of the study and designed the experiments. C.C., E.B., A.D., G.M., O.C., C.V.P., J.H., and R.B. collected samples. L.T. coordinated sample collection and performed the infection experiments. M.M. extracted the RNA and prepared the sequencing libraries. J.D.B. analyzed the results. L.B.B. and Y.G. supervised the project. J.D.B. wrote the paper with input from Y.G., L.T., L.B.B, and R.B.

## Supplementary data

### Supplementary Data S1

Supplementary Data S1 contains information on the 50 samples. Most variables describe the batch processing steps outlined in Supplementary Fig. S1. “id” is a unique identifier for each sample, “individual” is the individual identifier (“s” = susceptible, “r” = resistant), “status” is the susceptibility status, “treatment” is if the sample was infected or non-infected, “infection” is the date of the infection experiment (12 total), “arrival” is the identifier for the arrival batch (4 total), “extraction” is the batch for RNA extraction (5 total), “master mix” is the batch for library preparation (3 total), “rin” is the RNA Integrity Number from the Agilent Bioanalyzer, and “outlier” is a Boolean variable indicating if the sample was identified as an outlier (Supplementary Fig. S5) and removed from the analysis. (tds)

### Supplementary Data S2

Supplementary Data S2 contains the gene expression counts for the 11,336 genes after filtering lowly expressed genes for all 50 samples (Supplementary Fig. S2). Each row is a gene labeled with its Ensembl gene ID. Each column is a sample. Each sample is labeled according to the pattern “x##-status-treatment”, where x is “r” for resistant or “s” for susceptible, ## is the ID number, status is “resist” for resistant or “suscep” for susceptible, and treatment is “noninf” for non-infected or “infect” for infected. (tds)

### Supplementary Data S3

Supplementary Data S3 contains the results of the differential expression analysis with limma (Fig. 1). The workbook contains 4 sheets corresponding to the 4 tests performed. “status ni” is the test between resistant and susceptible individuals in the non-infected state, “status ii” is the test between resistant and susceptible individuals in the infected state, “treat resist” is the test between the non-infected and infected states for resistant individuals, and “treat_suscep” is the test between the non-infected and infected states for susceptible individuals. Each sheet has the same columns. “id” is the Ensembl gene ID, “gene” is the gene name, “logFC” is the log fold change from limma, “AveExpr” is the average log expression from limma, “t” is the t-statistic from limma, “P.Value” is the p-value from limma, “adj.P.Val” is the adjusted p-value from limma, “qvalue” is the q-value calculated with adaptive shrinkage, “chr” is the chromosome where the gene is located, “description” is the description of the gene from Ensembl, “phenotype” is the associated phenotype(s) assigned my Ensembl, “go_id” is the associated GO term(s) assigned by Ensembl, and “go_description” is the corresponding name(s) of the GO term(s). (xlsx)

### Supplementary Data S4

Supplementary Data S4 contains the results of the GWAS comparison analysis (Fig. 2). The first sheet “input-data” contains the p-values for the GWAS SNP assigned to each gene from each study. The columns “gwas_p_russia”, “gwas_p_gambia”, “gwas_p_ghana”, “gwas_p_uganda”, “gwas_p_height” contain the p-values from the TB susceptibility GWAS in Russia, The Gambia, Ghana, Uganda and Tanzania, and the height GWAS in Europeans, respectively. The columns “status_ni”, “status_ii”, “treat_resist”, and “treat_suscep” refer to the tests described for Supplementary Data S3 and contain the log fold changes for each comparison. All the other gene annotation columns are the same as described for Supplementary Data S3. The second sheet “top-genes” contains a subset of the full results to highlight those genes which had an absolute log fold change greater than 2 between resistant and susceptible individuals in the non-infected state (“status_ni”). (xlsx)

### Supplementary Data S5

Supplementary Data S5 contains the results of the classifier analysis. Specifically it contains the results from the support vector machine using the genes with a q-value less than 0.05 (Fig. 3). The sheet “gene-list” contains information about the genes used for the classifier (the columns are described in the section for Supplementary Data S3). The sheet “training-input” contains the input gene expression data for training the model. The sheet “training-results” contains the results of the leave-one-out-cross-validation when training the model on the samples from the current study. The sheet “testing-input” contains the input gene expression data for testing the model. The sheet “testing-results” contains the results from testing the model on the samples from Barreiro et al., 2012^24^. The column “prob_tb_suscep” is the probability of being susceptible to TB assigned by the model. (xlsx)

## Supplementary information

### Supplementary figures

**Figure S1.**
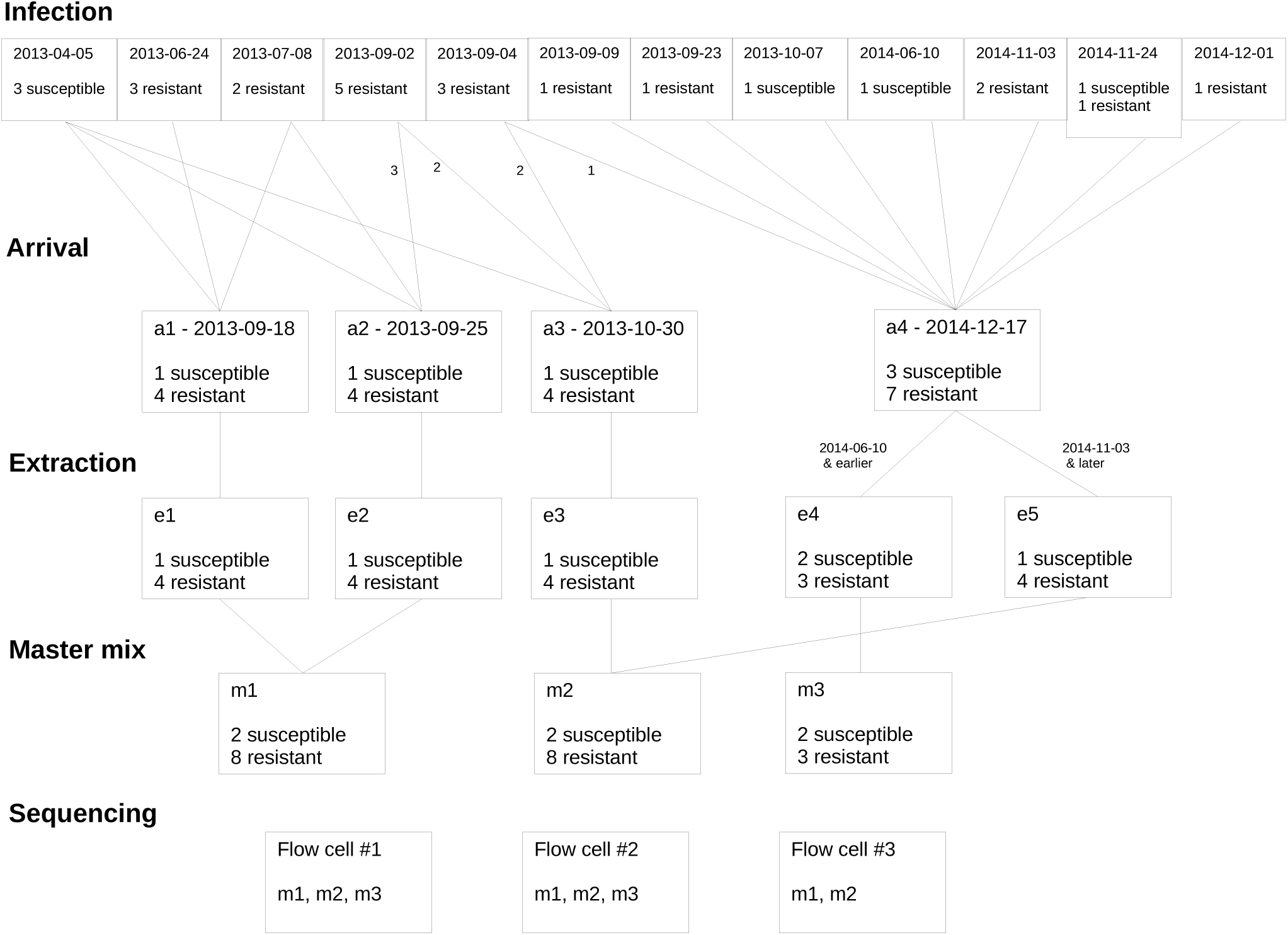
Batch processing. We designed the processing of the samples to minimize the introduction of technical batch effects. Specifically, we attempted to balance the processing of samples obtained from susceptible and resistant individuals. In the diagram, each box represents a batch. “Infection” labels the batches of the infection experiments, “Arrival” labels the batch shipments of cell lysates arrived in Chicago, USA from Paris, France, “Extraction” labels the batches of RNA extraction, “Master Mix” labels the batches of library preparation, and “Sequencing” labels the batches of flow cells. Each master mix listed in a flow cell batch was sequenced on only one lane of that flow cell.

**Figure S2.**
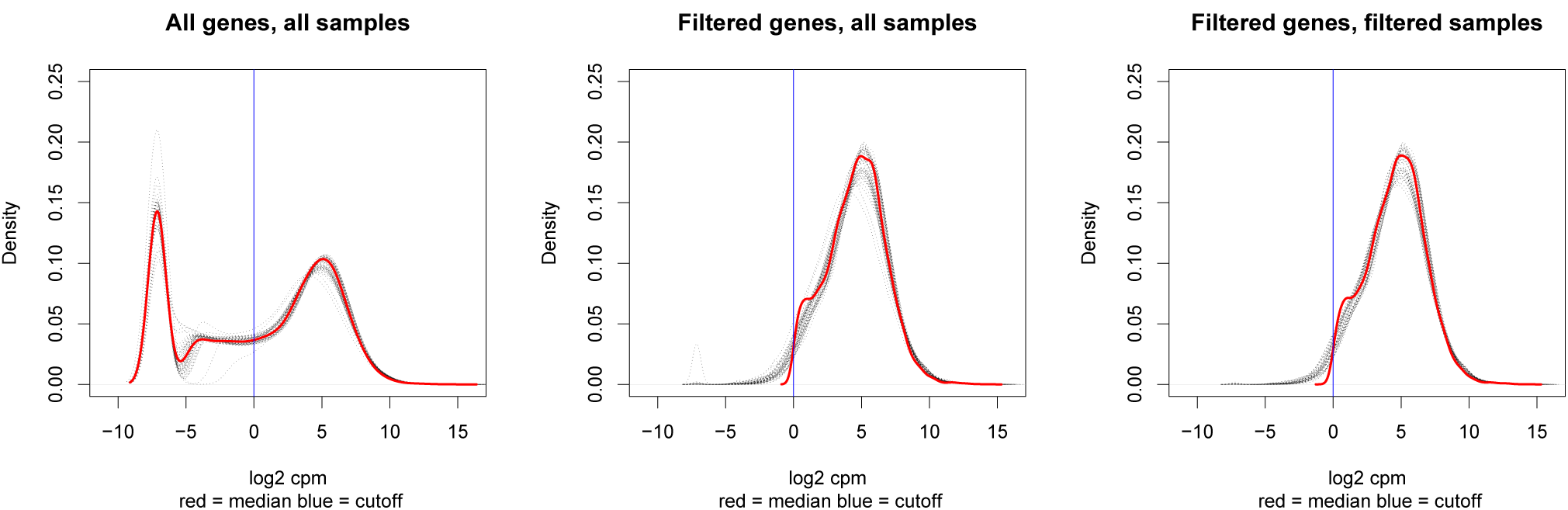
Gene expression distributions before and after filtering genes and samples. The log_2_ counts per million (cpm) of each sample is plotted as a dashed gray line. The solid red line represents the median value across all the samples. The vertical solid blue line at *x* = 0 represents the cutoff used to filter lowly expressed genes based on their median log_2_ cpm. The left panel is the data from all 19,800 genes and 50 samples, the middle panel is the data from the 11,336 genes remaining after removing lowly expressed genes, and the right panel is the data from 11,336 genes and the 44 samples remaining after removing outliers.

**Figure S3.**
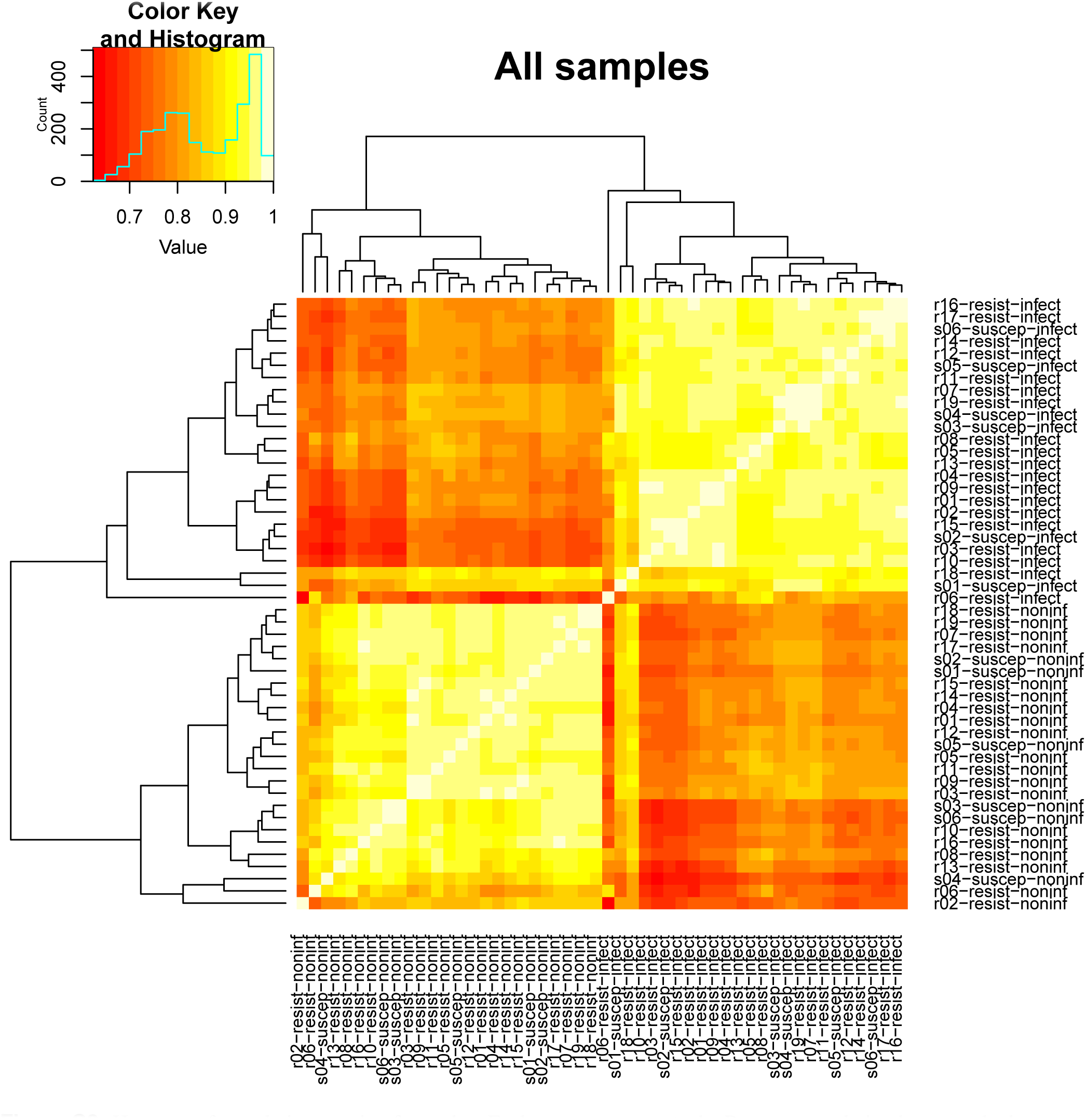
Heatmap of correlation matrix of samples. Each square represents the Pearson correlation between the log_2_ cpm expression values of two samples. Red indicates a low correlation of zero and white represents a high correlation of 1. The dendrogram displays the results of hierarchical clustering with the complete linkage method. The outliers of the non-infected samples are s04-suscept-noninf, r02-resist-noninf, and r06-resist-noninf. The outliers of the infected samples are s01-suscep-infect, r06-resist-infect, and r18-resist-infect.

**Figure S4.**
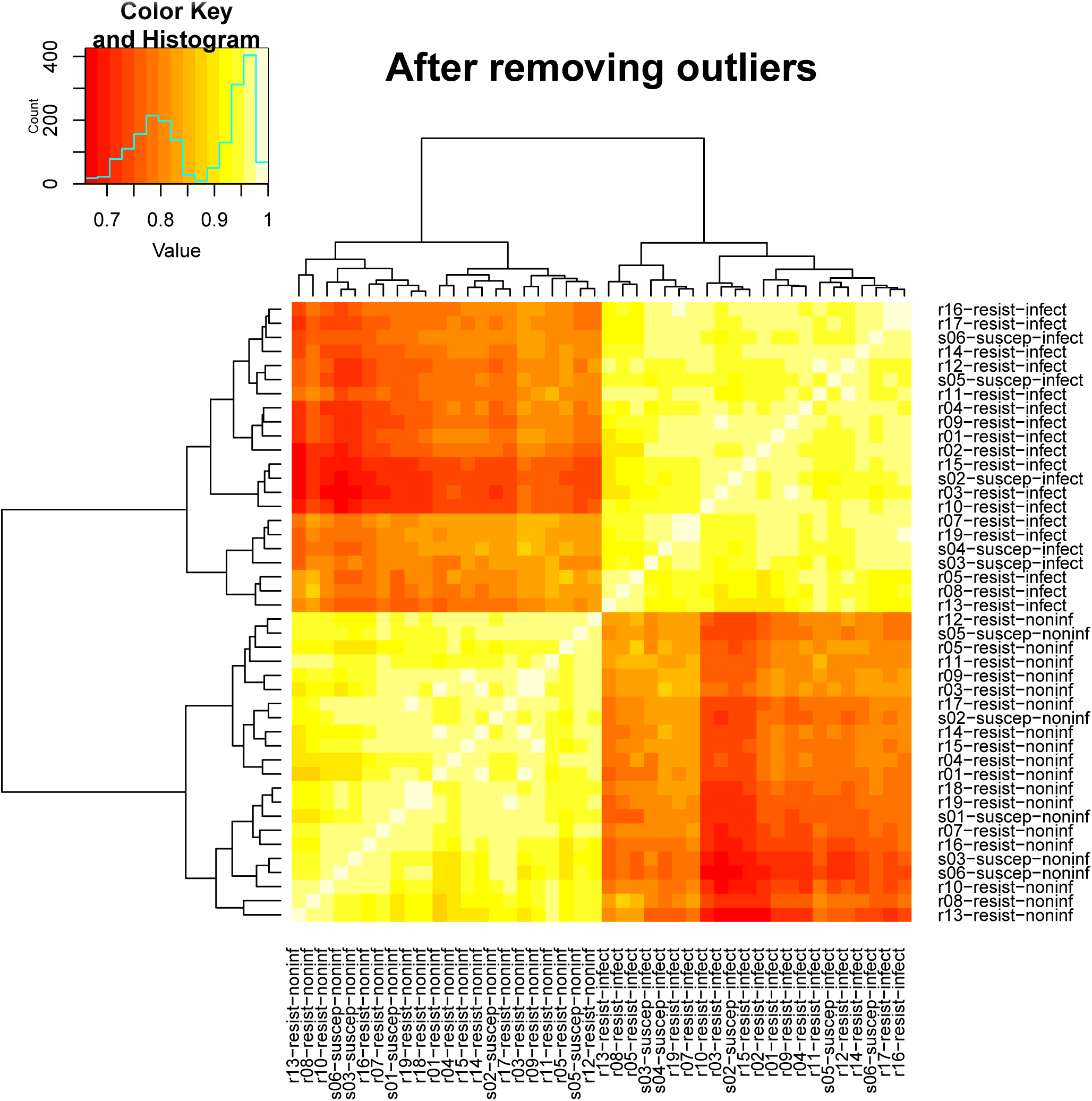
Heatmap of correlation matrix after removing outliers. Each square represents the Pearson correlation between the log_2_ cpm expression values of two samples. Red indicates a low correlation of zero and white represents a high correlation of 1. The dendrogram displays the results of hierarchical clustering with the complete linkage method.

**Figure S5.**
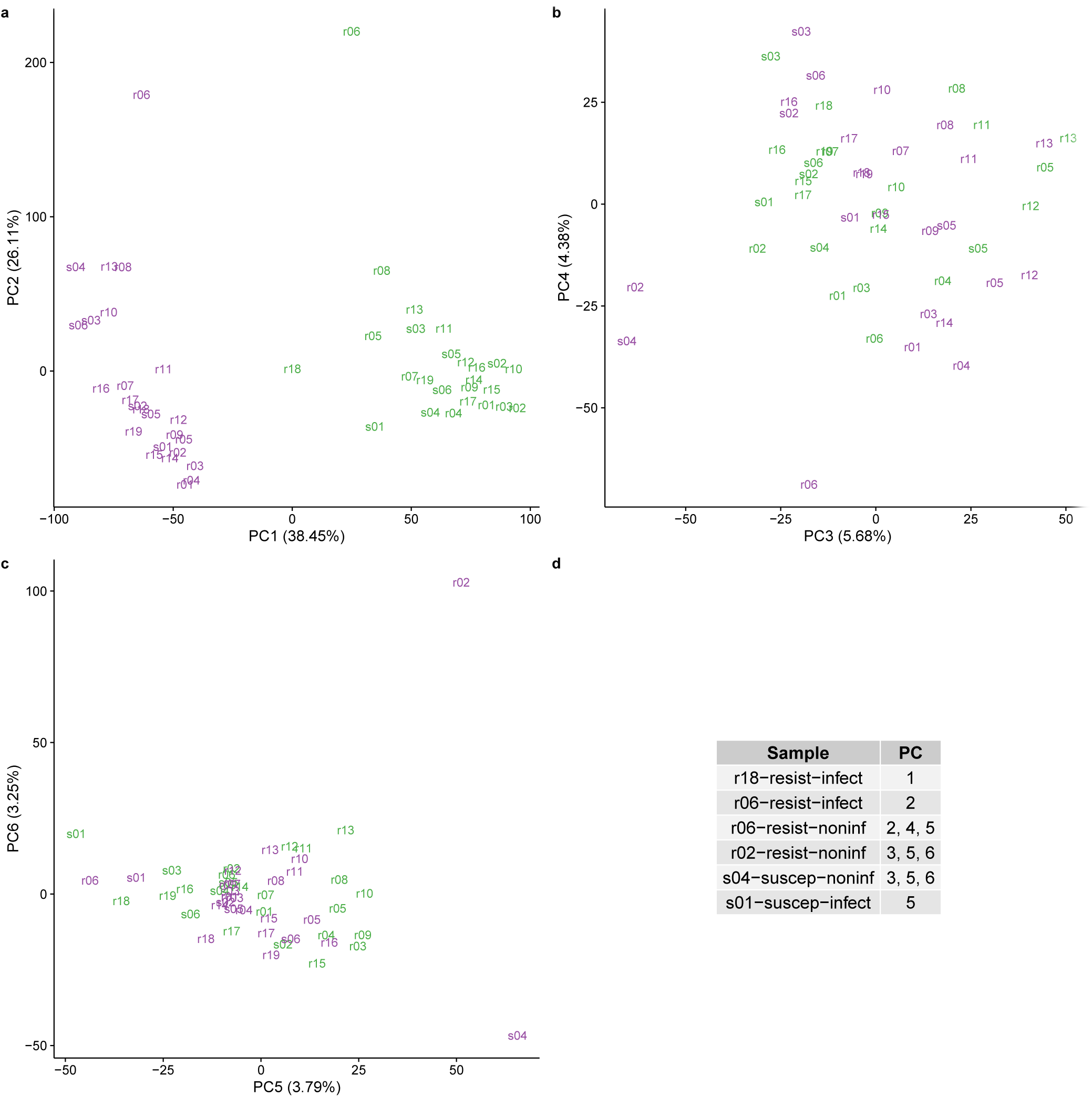
Principal components analysis (PCA) to identify outliers. (a) PC1 versus PC2, (b) PC3 versus PC4, and (c) PC5 versus PC6. Each sample is represented by its 3-letter ID. “s” stands for susceptible and “r” for resistant, and the text is colored on the basis of treatment status (purple is non-infected; green is infected). The value in parentheses in each axis is the percentage of total variation accounted for by that PC. The outliers are listed in (d). These samples do not fall within 2 standard deviations of the mean value of the PCs listed in the right column. Note that a separate mean was calculated for the non-infected and infected samples for PC1 only.

**Figure S6.**
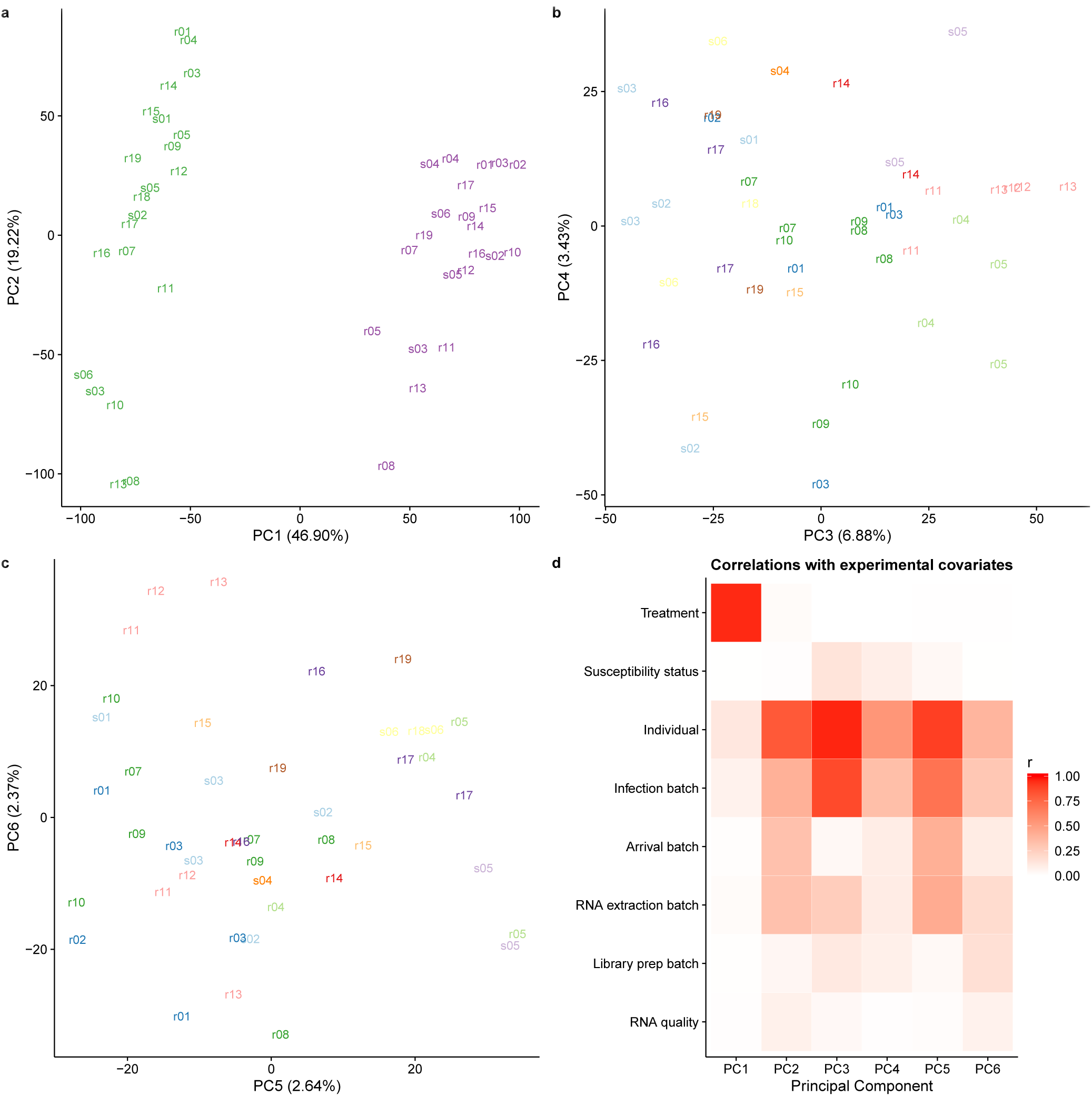
Check for technical batch effects using principal components analysis (PCA). (a) PC1 versus PC2. The text labels are the individual identifiers. Purple indicates non-infected samples and green indicates infected. (b) PC3 versus PC4. The colors indicate the different infection batches. (c) PC5 versus PC6. The colors indicate the different infection batches. (d) The Pearson correlation of PCs 1-6 with each of the recorded biological and technical covariates. The correlations vary from 0 (white) to 1 (red).

**Figure S7.**
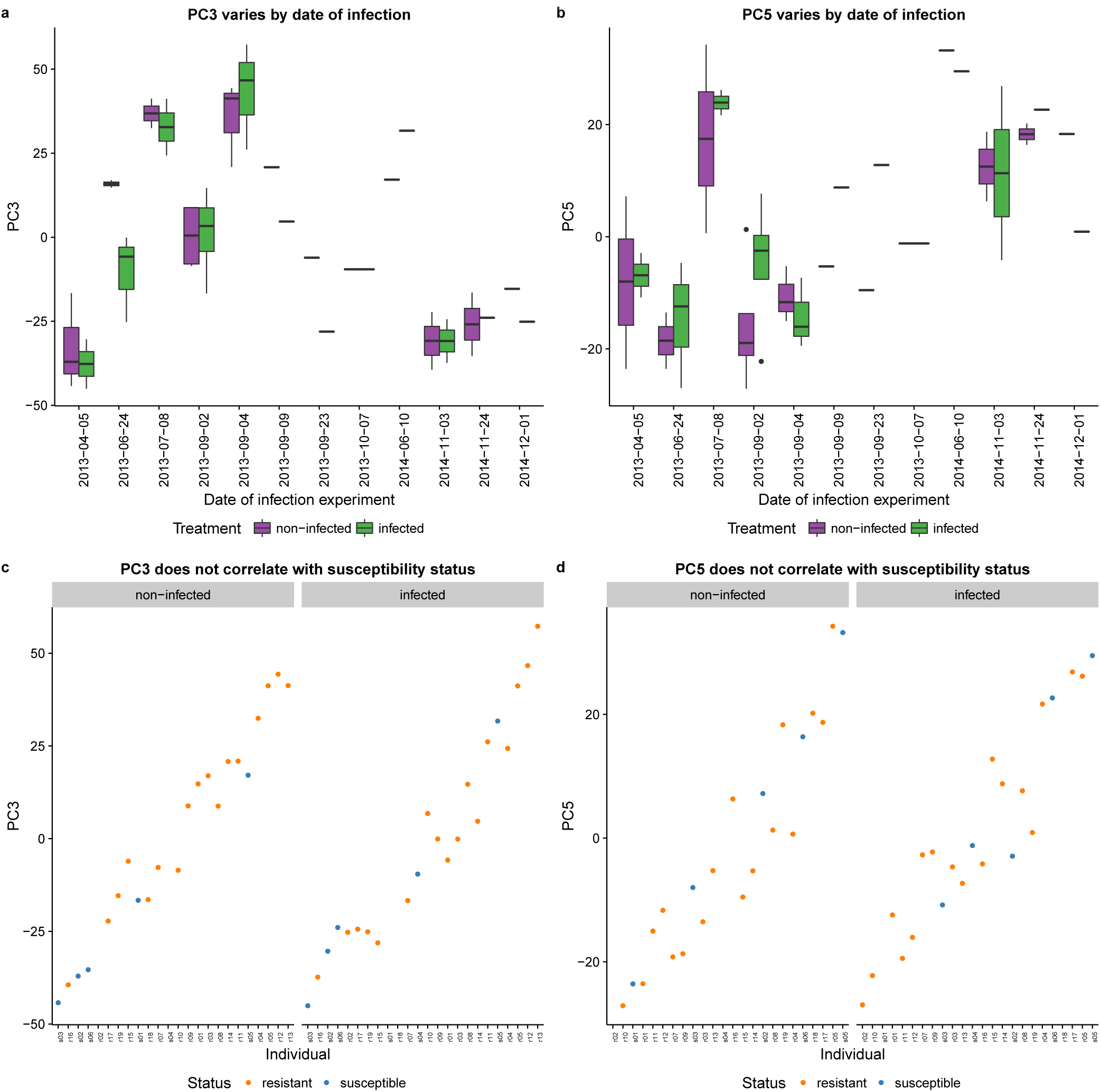
Check for confounding effect of infection batch. PC3 (a) and PC5 (b) varied by the date of infection. Non-infected samples are in purple and infected samples in green. Importantly, however, this technical variation arising from infection batch did not correlate with the susceptibility status of the individuals (c and d). Resistant individuals are in orange and susceptible individuals in blue.

**Figure S8.**
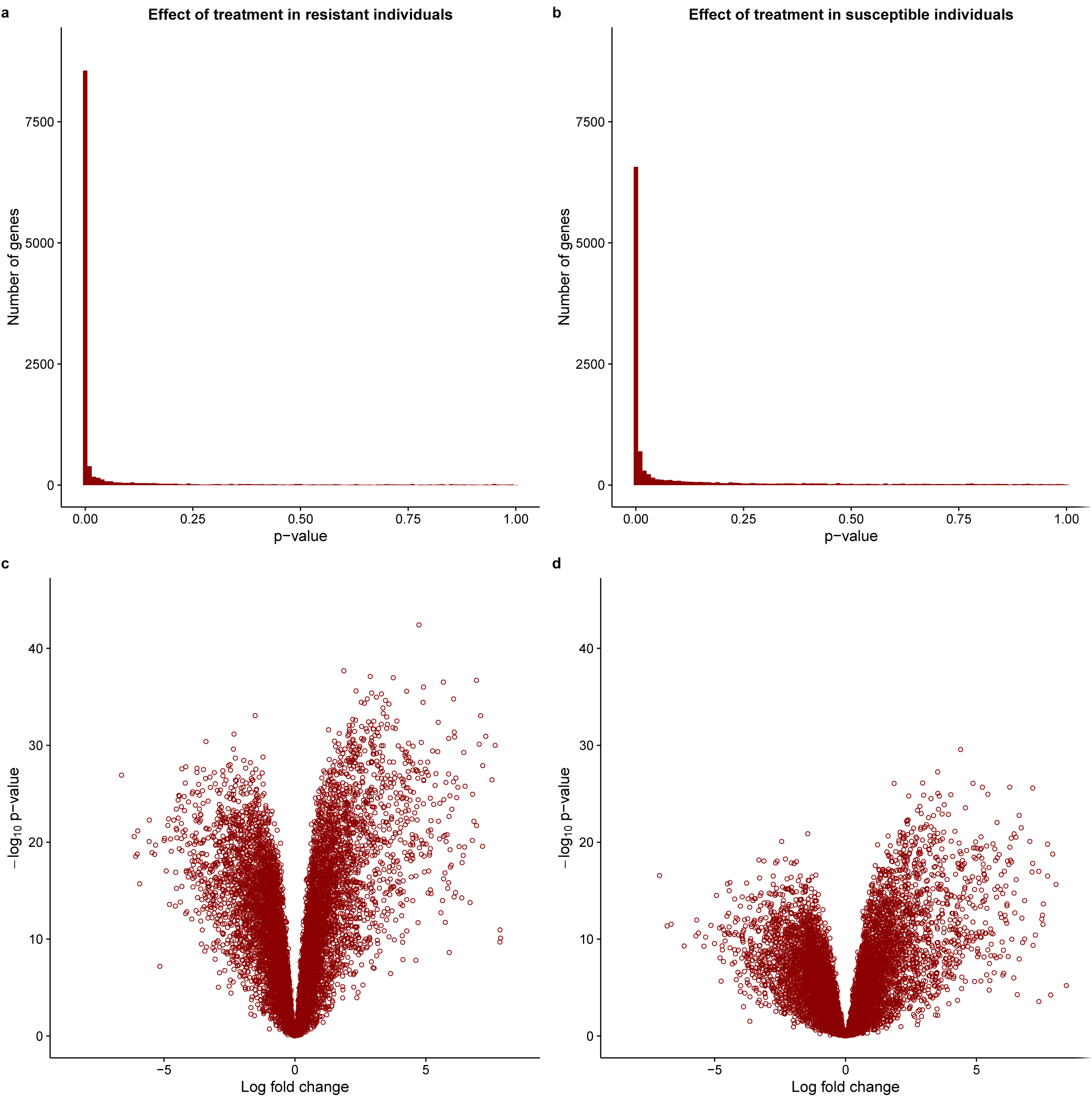
Effect of treatment with MTB. The top panel contains the distribution of unadjusted p-values after testing for differential expression between the non-infected and infected states in (a) resistant and (b) susceptible individuals. The bottom panel contains the corresponding volcano plots for the (c) resistant and (d) susceptible individuals. The x-axis is the log fold change in gene expression level between susceptible and resistant individuals and the y-axis is the –log_10_ p-value. Red indicates genes which are significant differentially expressed with a q-value less than 10%. Because of the extremely skewed p-value distribution, all genes are significantly differentially expressed at this false discovery rate.

**Figure S9.**
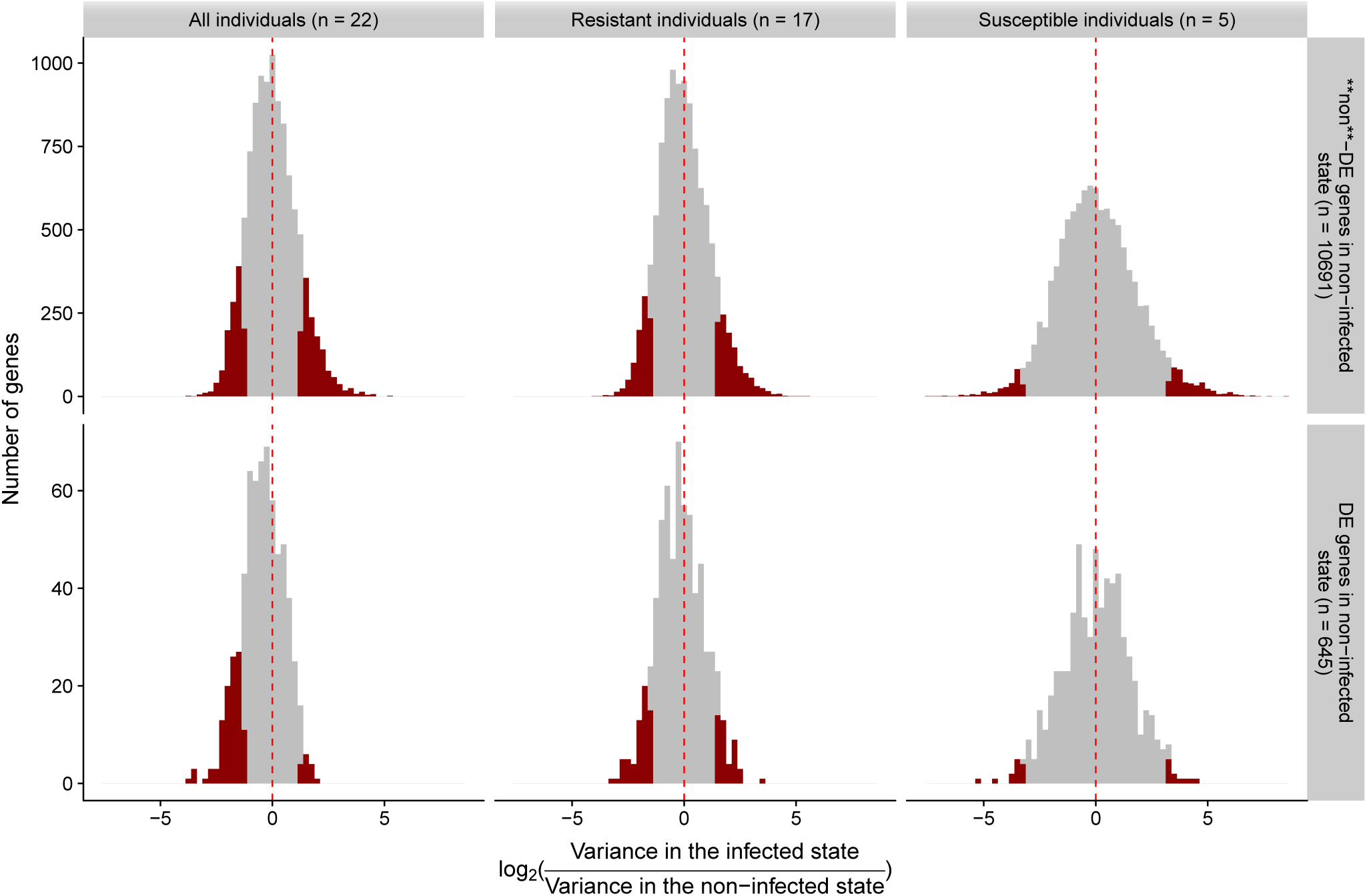
Check for systematic differences in gene expression variance between the infected and non-infected states. We identified DE genes between susceptible and resistant individuals in the non-infected but not the infected state. Could this be a statistical artifact due to an overall increase in gene expression variance upon infection thus reducing power to detect DE genes? No, because we did not observe an overall increase in gene expression variance in the infected state. The histograms show the distribution of the log_2_-transformed ratio of the gene expression variance in the infected state to the variance in the non-infected state. If there was an overall increase in variance, the distributions should be shifted towards the right, but instead they are all symmetrical. The top row shows the results for the 10,691 genes which were not differentially expressed between susceptible and resistant individuals in the non-infected state, and the bottom row shows the results for the 645 genes which were. The left column shows the results for all 22 individuals in the study, the middle column for the 17 resistant individuals, and the right column for the 5 susceptible individuals (note that the right column has the widest spread because of this small sample size). Highlighted in red are genes which had a *P* < 0.05 from an F test comparing the two variances. The number of genes with a significant increase or decrease in variance was also mostly symmetrical (decrease vs. increase starting at top left panel and proceeding clockwise: 1,232 vs. 1,362; 934 vs. 1,118; 275 vs. 455; 13 vs. 11; 64 vs. 44; 108 vs. 15).

**Figure S10.**
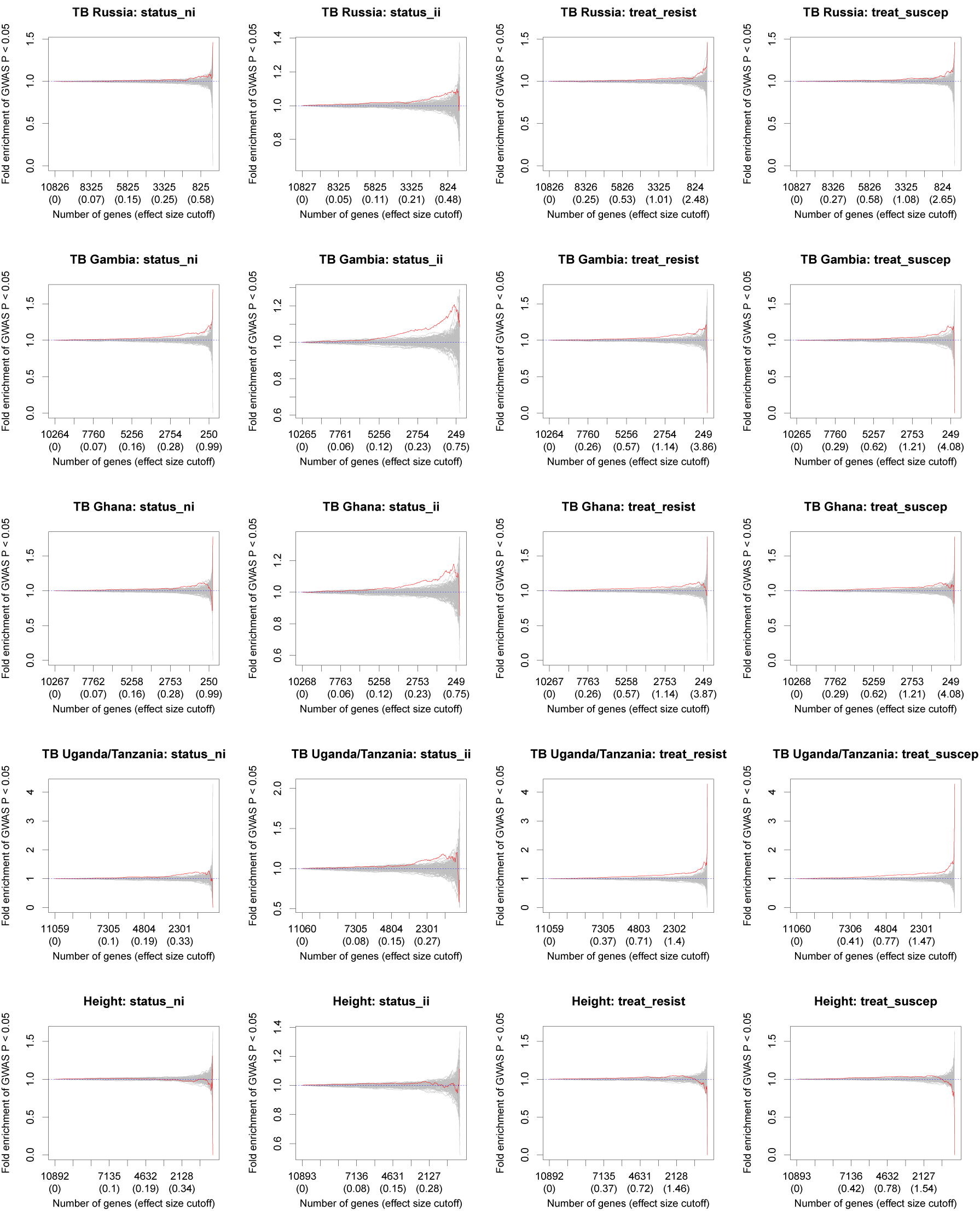
Comparison of differential expression and GWAS results. In each subplot, the y-axis is the fold enrichment (y-axis) of genes assigned a SNP with p-value less than 0.05 from the GWAS. The x-axis is bins of genes with increasingly stringent effect size cutoffs of the absolute log fold change for the different expression contrast. The effect size cutoffs were chosen such that each bin from left to right contained approximately 25 fewer genes. The red line is the results from the actual data. The grey lines are the results from 100 permutations. The dashed blue line at y=1 is the null expectation. The rows correspond to the 5 GWAS studies: Russia^18^, The Gambia^13^, Ghana^13^, Uganda and Tanzania^19^, and height in individuals of European ancestry^25^. The columns correspond to the 4 differential expression contrasts: resistant vs. susceptible individuals in the non-infected state (status ni), resistant vs. susceptible individuals in the infected state (status ii), effect of treatment in resistant individuals (treat resist), and effect of treatment in susceptible individuals (treat suscep). The x-axis slightly varies based on the number of genes that were able to be assigned a nearby SNP for each GWAS, and thus is consistent only within each study (i.e. row, although the exact tick labels in each plot slightly vary based on R’s rules for annotating axes). The y-axis is set separately for each plot based on the minimum and maximum fold enrichment values for that particular analysis.

**Figure S11.**
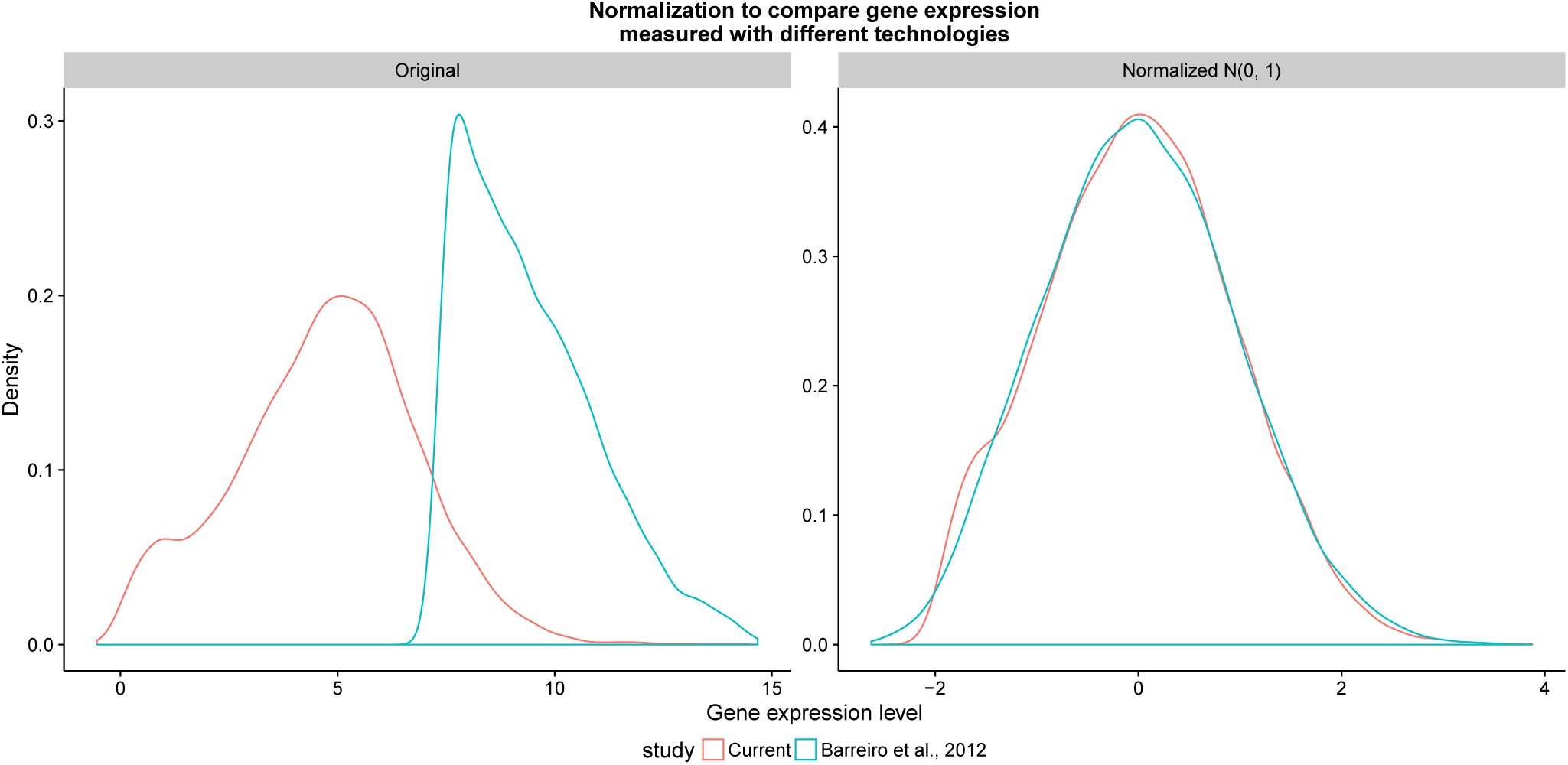
Normalizing gene expression distributions. (left) The distribution of the median log_2_ cpm of the RNA-seq data from the current study in red compared to the distribution of the median gene expression levels of the microarray data from Barreiro et al., 2012^24^ in blue. (right) The distributions of the same data sets after normalizing each sample to a standard normal distribution.

**Figure S12.**
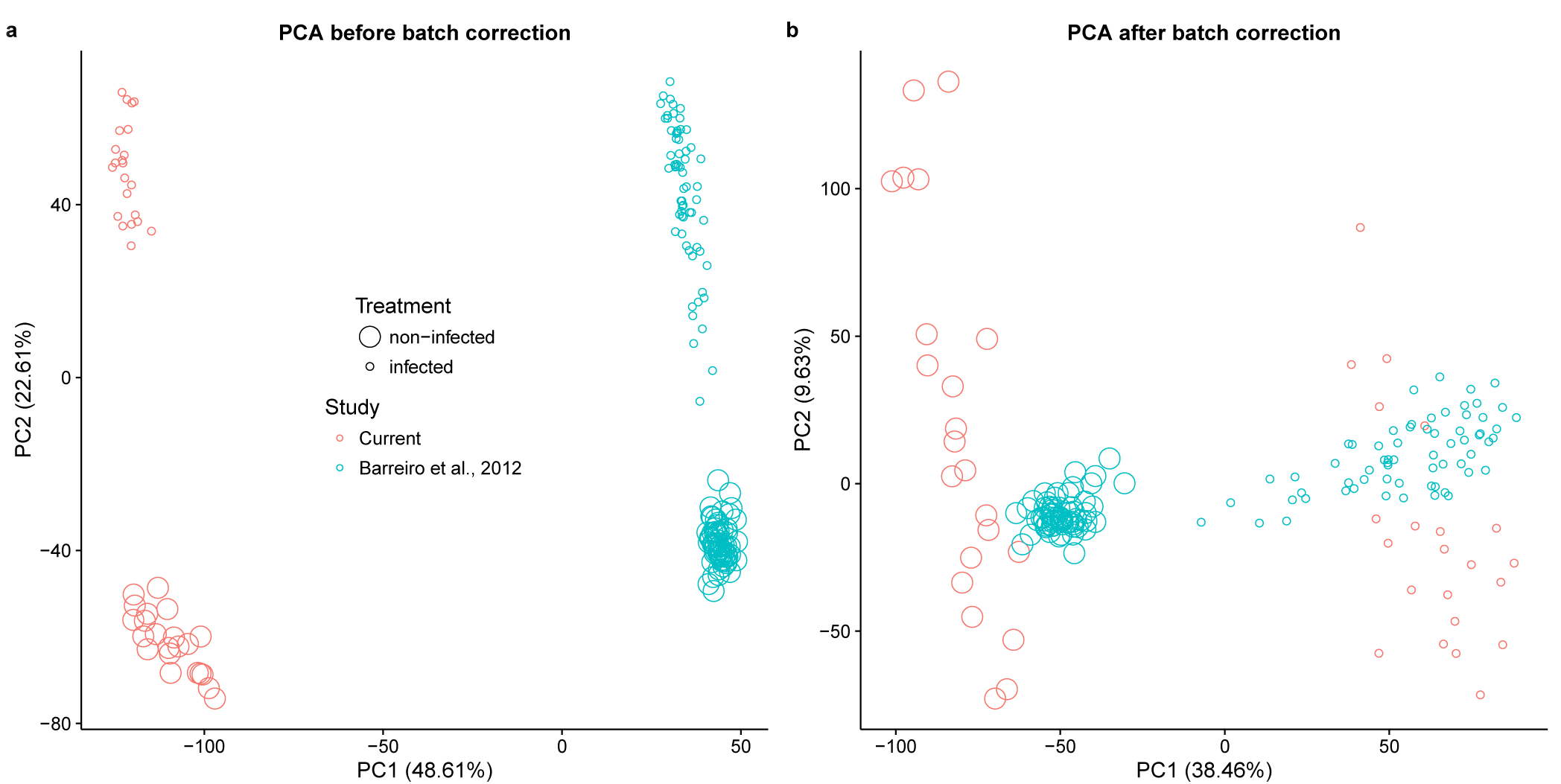
Principal components analysis (PCA) of combined data sets. (a) PC1 versus PC2 of the combined data set of the RNA-seq data from the current study (red) and the microarray data from Barreiro et al., 2012^24^ (blue). The large circles are non-infected samples, and the small circles are infected samples. The value in parentheses is the percentage of the total variation accounted for by that PC. (b) The same data after regressing the original PC1 in (a).

**Figure S13.**
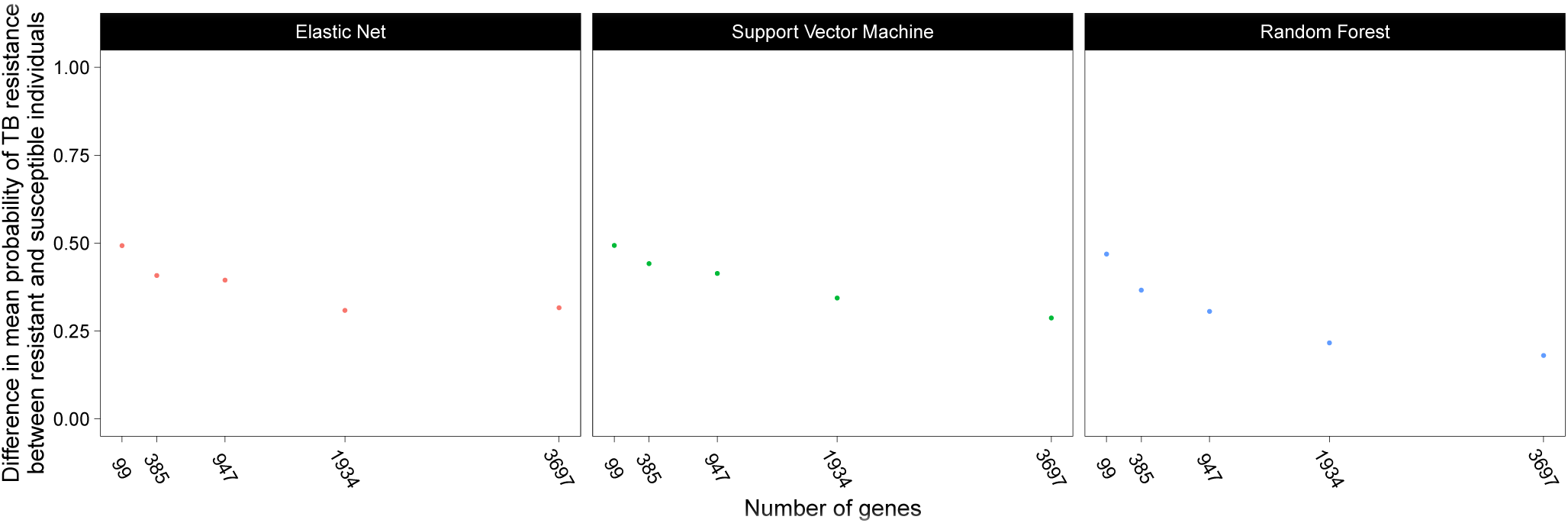
Comparing the classification results of different methods and number of input genes. We compared 3 different machine learning methods (elastic net, support vector machine, random forest) and used 5 different sets of input genes. The input genes (x-axis) were obtained by varying the q-value cutoff for differential expression between susceptible and resistant individuals in the non-infected state from 5% to 25%. The evaluation metric (y-axis) was the difference of the mean assigned probability of being TB resistant between the known resistant and susceptible individuals in the current study.

**Figure S14.**
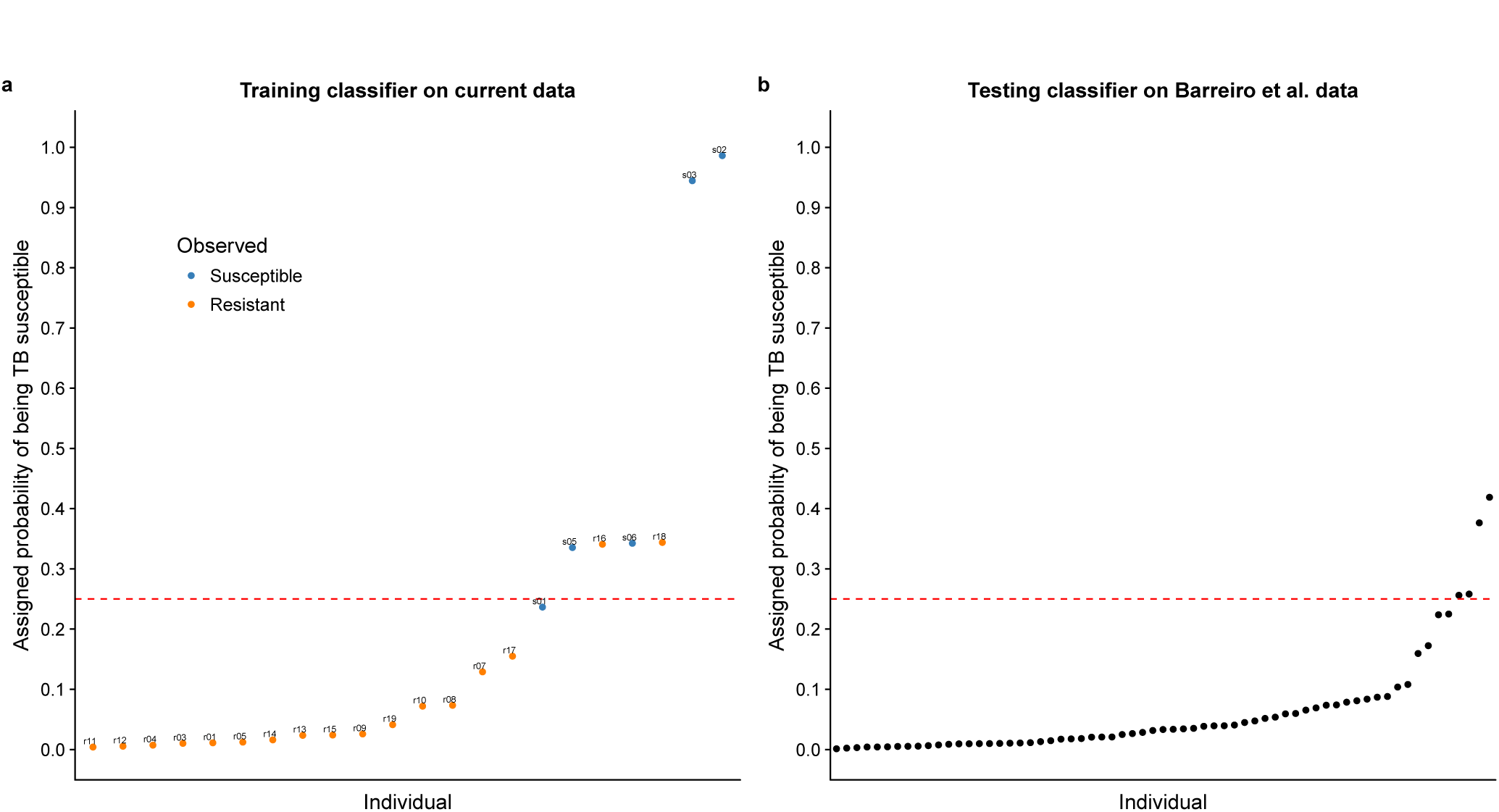
Classifying TB susceptible individuals using an elastic net model. (a) The estimates of predicted probability of TB susceptibility from the leave-one-out-cross-validation for individuals in the current study. The blue circles represent individuals known to be susceptible to TB, and orange those resistant to TB. The horizontal blue line at a probability of 0.25 almost separates susceptible and resistant individuals. (b) The estimates of predicted probability of TB susceptibility from applying the classifier trained on the data from the current study to a test set of independently collected healthy individuals^24^.

**Figure S15.**
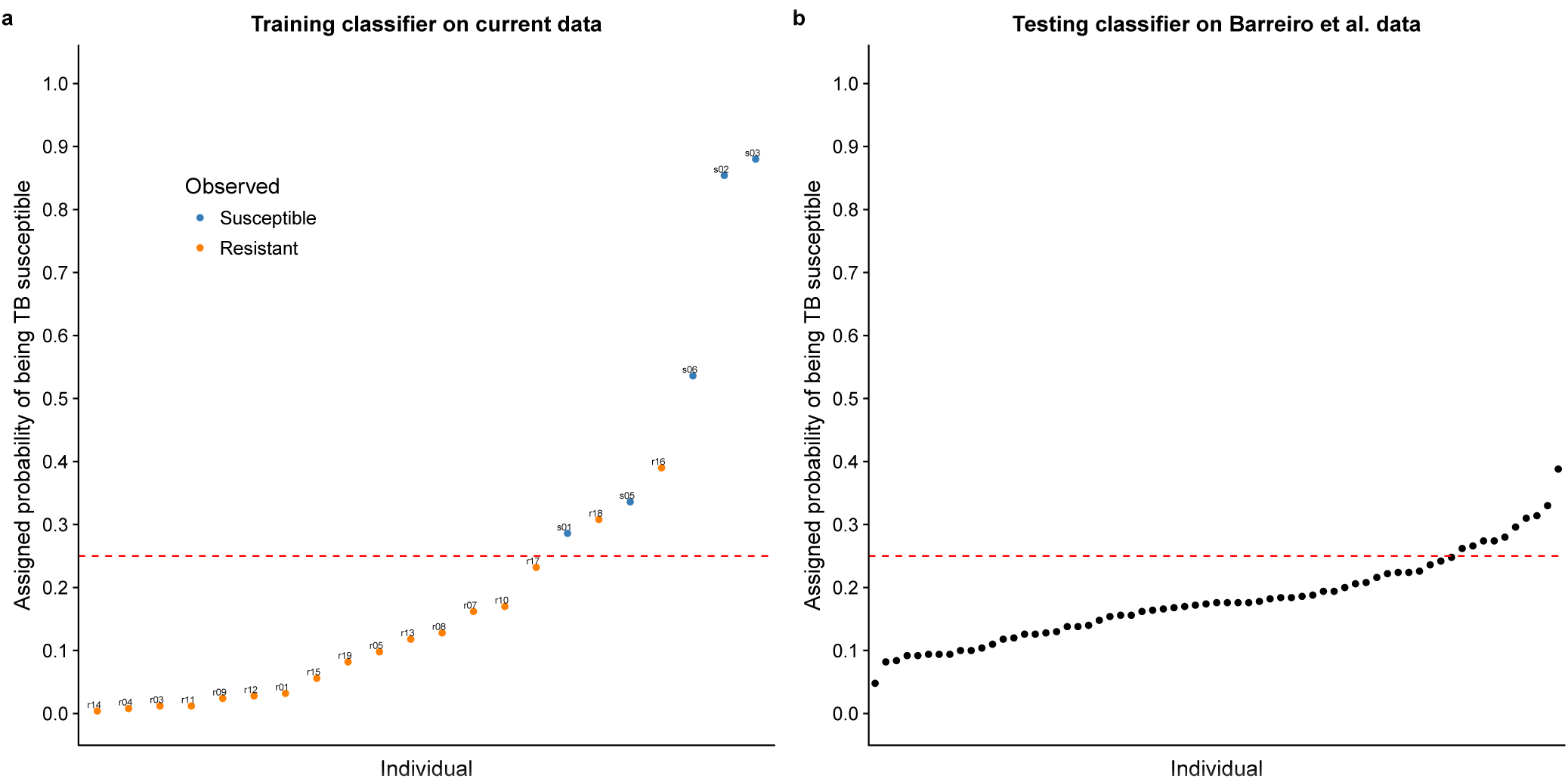
Classifying TB susceptible individuals using a random forest model. (a) The estimates of predicted probability of TB susceptibility from the leave-one-out-cross-validation for individuals in the current study. The blue circles represent individuals known to be susceptible to TB, and orange those resistant to TB. The horizontal blue line at a probability of 0.25 separates susceptible and resistant individuals. (b) The estimates of predicted probability of TB susceptibility from applying the classifier trained on the data from the current study to a test set of independently collected healthy individuals^24^.

**Figure S16.**
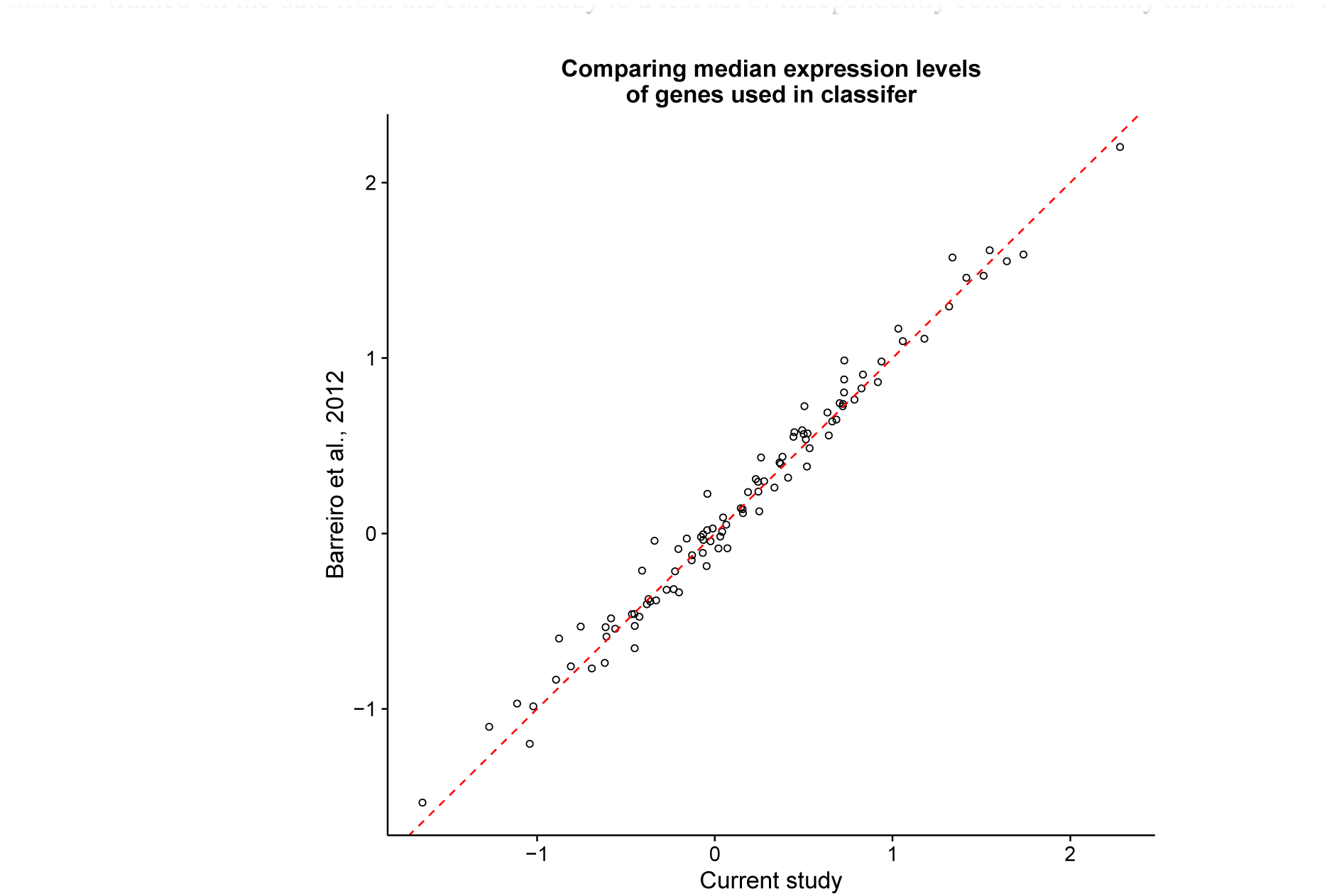
Comparing gene expression between the two studies. After normalization and batch-correction, the median expression levels of the 99 genes used in the classifier were similar between the samples in the current study and those in Barreiro et al., 2012^24^. The dashed red line is the 1:1 line.

**Figure S17.**
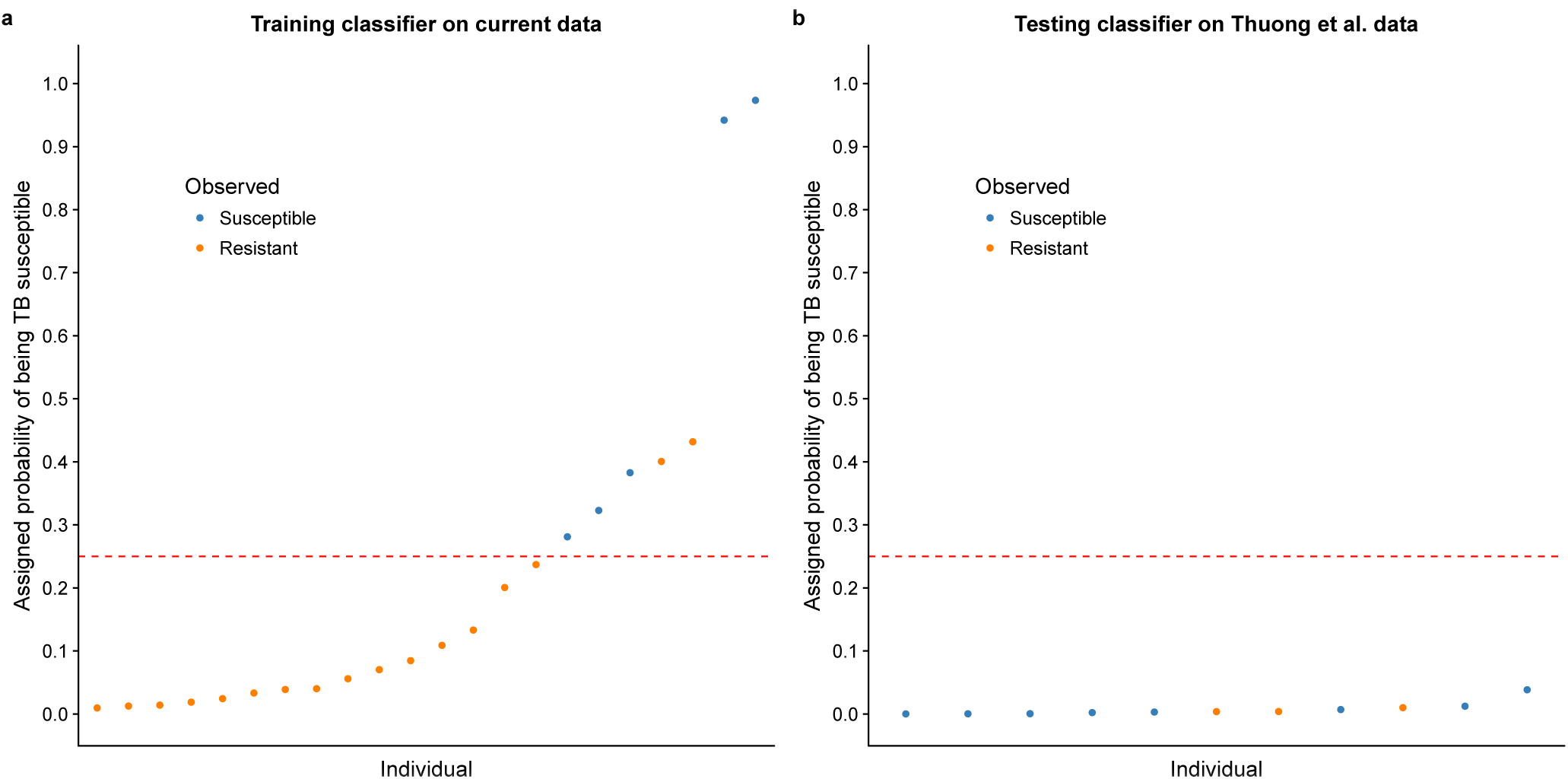
Classifying individuals from Thuong et al., 2008^20^ using a support vector machine model. We followed the same training and testing procedure performed for testing the classifier described in the main text (Fig. 3, see Classifier in Methods). Not surprisingly since the data sets were from different cell types, the classifier trained on the dendritic cells in this study performed poorly when tested on samples with gene expression levels measured in macrophages. To match our naming system, we labeled the individuals from Thuong et al., 2008^20^ with latent TB as resistant (n = 3 after removing the outlier sample LTB2) and the individuals recovered from pulmonary or meningeal TB as susceptible (n = 4 each). (a) The estimates of predicted probability of TB susceptibility from the leave-one-out-cross-validation for individuals in the current study. The blue circles represent individuals known to be susceptible to TB, and orange those resistant to TB. The horizontal dashed red line at a probability of 0.25 separates susceptible and resistant individuals. (b) The estimates of predicted probability of TB susceptibility from applying the classifier trained on the data from the current study to a test set of putatively susceptible and resistant individuals^20^.

## References

World Health Organization. Global tuberculosis report 2015 (2015).

World Health Organization. Global TB facts 2015 (2015).

Glaziou, P., Sismanidis, C., Floyd, K. & Raviglione, M. Global epidemiology of tuberculosis. Cold Spring Harbor perspectives in medicine 5, 445–461 (2015).

Sotgiu, G., Centis, R., D’ambrosio, L. & Migliori, G. B. Tuberculosis treatment and drug regimens. Cold Spring Harbor perspectives in medicine 5, 505–516 (2015).

Seung, K. J., Keshavjee, S. Rich, M. L. Multidrug-resistant tuberculosis and extensively drug-resistant tuberculosis. Cold Spring Harbor perspectives in medicine 5, 579–598 (2015).

Muñoz,, L., Stagg, H. R., & Abubakar, I., Diagnosis and management of latent tuberculosis infection. Cold Spring Harbor perspectives in medicine 5, 517–529 (2015).

North, R. J. & Jung, Y.-J.Immunity to tuberculosis. Annual review of immunology 22, 599–623 (2004).

O’Garra, A. et al. The immune response in tuberculosis. Annual review of immunology 31, 475–527 (2013).

Kallmann, F. J. Reisner, D. Twin studies on genetic variations in resistance to tuberculosis. Journal of Heredity 34, 269–276 (1943).

Comstock, G. W. Tuberculosis in twins: a re-analysis of the Prophit survey. The American review of respiratory disease 117, 621–4 (1978).

Cobat, A. et al. High heritability of antimycobacterial immunity in an area of hyperendemicity for tuberculosis disease. The Journal of infectious diseases 201, 15–9 (2010).

Möller, M & Hoal E. G Current findings, challenges and novel approaches in human genetic susceptibility to tuberculosis. Tuberculosis (Edinburgh, Scotland) 90, 71–83 (2010).

Thye, T. et al. Genome-wide association analyses identifies a susceptibility locus for tuberculosis on chromosome 18q11.2. Nature genetics 42, 739–41 (2010).

Mahasirimongkol, S. et al. Genome-wide association studies of tuberculosis in Asians identify distinct at-risk locus for young tuberculosis. Journal of human genetics 57, 363–7 (2012).

Thye, T. et al. Common variants at 11p13 are associated with susceptibility to tuberculosis. Nature genetics 44, 257–9 (2012).

Png, E. et al. A genome wide association study of pulmonary tuberculosis susceptibility in Indonesians. BMC medical genetics 13, 1–9 (2012).

Chimusa, E. R. et al. Genome-wide association study of ancestry-specific TB risk in the South African Coloured population. Human molecular genetics 23, 796–809 (2014).

Curtis, J. et al. Susceptibility to tuberculosis is associated with variants in the ASAP1 gene encoding a regulator of dendritic cell migration. Nature genetics 47, 523–7 (2015).

Sobota, R. S. et al. A locus aprocessing. We designed the processing of the samples to minimizet 5q33.3 confers resistance to tuberculosis in highly susceptible individuals. American journal of human genetics 98, 514–24 (2016).

Thuong, N. T. T. et al. Identification of tuberculosis susceptibility genes with human macrophage gene expression profiles. PLoS pathogens 4, e1000229 (2008).

Bryant, P. A. et al. Susceptibility to acute rheumatic fever based on differential expression of genes involved in cytotoxicity, chemotaxis, and apoptosis. Infection and immunity 82, 753–61 (2014).

Loddenkemper, R., Lipman, M. & Zumla, A. Clinical aspects of adult tuberculosis. Cold Spring Harbor perspectives in medicine 6, a017848 (2016).

Cooper, A. M. Cell-mediated immune responses in tuberculosis. Annual review of immunology 27, 393–422 (2009).

Barreiro, L. B. et al. Deciphering the genetic architecture of variation in the immune response to Mycobacterium tuberculosis infection. Proceedings of the National Academy of Sciences of the United States of America 109, 1204–9 (2012)

Lango Allen, H. et al. Hundreds of variants clustered in genomic loci and biological pathways affect human height. Nature 467, 832–8 (2010).

Tailleux, L. et al. Probing host pathogen cross-talk by transcriptional profiling of both Mycobacterium tuberculosis and infected human dendritic cells and macrophages. PloS one 3, e1403 (2008).

Deretic, V. Autophagy in tuberculosis. Cold Spring Harbor Perspectives in Medicine 4, 53–67 (2014).

Castrejó-Jiménez,, N. S., Leyva-Paredes, K., Hernández-González,, J. C., Luna-Herrera, J. & García-Pérez,, B. E. The role of autophagy in bacterial infections. Bioscience trends 9, 149–59 (2015).

Spang, N. et al. RAB3GAP1 and RAB3GAP2 modulate basal and rapamycin-induced autophagy. Autophagy 10, 2297–309 (2014).

Sturgill-Koszycki, S. et al. Lack of acidification in mycobacterium phagosomes produced by exclusion of the vesicular proton-atpase. Science (New York, N.Y.) 263, 678–81 (1994).

Hornef, M. W., Wick, M. J., Rhen, M. & Normark, S. Bacterial strategies for overcoming host innate and adaptive immune responses. Nature immunology 3, 1033–40 (2002).

Hestvik, A. L. K., Hmama, Z. & Av-Gay, Y. Mycobacterial manipulation of the host cell. FEMS microbiology reviews 29, 1041–50 (2005).

Flynn, J. L., Goldstein, M. M., Triebold, K. J., Koller, B. & Bloom, B. R. Major histocompatibility complex class I-restricted T cells are required for resistance to Mycobacterium tuberculosis infection. Proceedings of the National Academy of Sciences of the United States of America 89, 12013–7 (1992).

Grotzke, J. E. et al. The Mycobacterium tuberculosis phagosome is a HLA-I processing competent organelle. PLoS pathogens 5, e1000374 (2009).

Grotzke, J. E., Siler, A. C., Lewinsohn, D. A. & Lewinsohn, D. M. Secreted immunodominant Mycobacterium tuberculosis antigens are processed by the cytosolic pathway. Journal of immunology (Baltimore, Md.: 1950) 185, 4336–43 (2010).

Lindestam Arlehamn, C. S., Lewinsohn, D., Sette, A. & Lewinsohn, D. Antigens for CD4 and CD8 T cells in tuberculosis. Cold Spring Harbor perspectives in medicine 4, 89–103 (2014).

Miller, M. D. & Krangel, M. S. The human cytokine I-309 is a monocyte chemoattractant. Proceedings of the National Academy of Sciences of the United States of America 89, 2950–4 (1992).

Blischak, J. D., Tailleux, L., Mitrano, A., Barreiro, L. B. & Gilad, Y. Mycobacterial infection induces a specific human innate immune response. Scientific reports 5, 16882 (2015).

Sudhof, T. C. The synaptic vesicle cycle. Annual review of neuroscience 27, 509–47 (2004).

Berry, M. P. R. et al. An interferon-inducible neutrophil-driven blood transcriptional signature in human tuberculosis. Nature 466, 973–7 (2010).

Blankley, S. et al. The application of transcriptional blood signatures to enhance our understanding of the host response to infection: the example of tuberculosis. Philosophical transactions of the Royal Society of London. Series B, Biological sciences 369, 20130427 (2014).

Maertzdorf, J., Kaufmann, S. H. & Weiner, J. Toward a unified biosignature for tuberculosis. Cold Spring Harbor Perspectives in Medicine 5, 183–95 (2015).

Chaussabel, D. et al. Unique gene expression profiles of human macrophages and dendritic cells to phylogenetically distinct parasites. Blood 102, 672–81 (2003).

Liao, Y., Smyth, G. K. & Shi, W. The Subread aligner: fast, accurate and scalable read mapping by seed-and-vote. Nucleic acids research 41, e108 (2013).

Yates, A. et al. Ensembl 2016. Nucleic acids research 44, D710–6 (2016).

Huber, W. et al. Orchestrating high-throughput genomic analysis with Bioconductor. Nature methods 12, 115–21 (2015).

Durinck, S. et al. BioMart and Bioconductor: a powerful link between biological databases and microarray data analysis. Bioinformatics (Oxford, England) 21, 3439–40 (2005).

Durinck, S., Spellman, P. T., Birney, E. & Huber, W. Mapping identifiers for the integration of genomic datasets with the R/Bioconductor package biomaRt. Nature protocols 4, 1184–91 (2009).

Liao, Y., Smyth, G. K. & Shi, W. featureCounts: an efficient general purpose program for assigning sequence reads to genomic features. Bioinformatics (Oxford, England) 30, 923–30 (2014).

Smyth, G. K. Linear models and empirical bayes methods for assessing differential expression in microarray experiments. Statistical applications in genetics and molecular biology 3, Article3 (2004).

Law, C. W., Chen, Y., Shi, W. & Smyth, G. K. voom: Precision weights unlock linear model analysis tools for RNA-seq read counts. Genome biology 15, R29 (2014).

Ritchie, M. E. et al. limma powers differential expression analyses for RNA-sequencing and microarray studies. Nucleic acids research 43, e47 (2015).

Smyth, G. K., Michaud, J. & Scott, H. S. Use of within-array replicate spots for assessing differential expression in microarray experiments. Bioinformatics (Oxford, England) 21, 2067–75 (2005).

Liu, R. et al. Why weight? modelling sample and observational level variability improves power in RNA-seq analyses. Nucleic acids research 43, e97 (2015).

Stephens, M. False discovery rates: a new deal. Biostatistics (Oxford, England) 1–20 (2016).

Lawrence, M. et al. Software for computing and annotating genomic ranges. PLoS computational biology 9, e1003118 (2013).

Jurasinski, G., Koebsch, F., Guenther, A. & Beetz, S. flux: Flux rate calculation from dynamic closed chamber measurements (2014). R package version 0.3–0.

Kuhn, M. Building predictive models in R using the caret package. Journal of Statistical Software 28, 1–26 (2008).

Friedman, J., Hastie, T. & Tibshirani, R. Regularization paths for generalized linear models via coordinate descent. Journal of Statistical Software 33, 2008–2010 (2010).

Karatzoglou, A., Smola, A., Hornik, K. & Zeileis, A. kernlab - an S4 package for kernel methods in R. Journal of Statistical Software 11, 1–20 (2004).

Liaw, A. & Wiener, M. Classification and regression by randomForest. R News 2, 18–22 (2002).

Köster,, J. & Rahmann, S. Snakemake–a scalable bioinformatics workflow engine. Bioinformatics (Oxford, England) 28, 2520–2 (2012).

Ewels, P., Magnusson, M., Lundin, S. & Käller,, M. MultiQC: summarize analysis results for multiple tools and samples in a single report. Bioinformatics (Oxford, England) btw354 (2016).

Li, H. et al. The sequence alignment/map format and samtools. Bioinformatics (Oxford, England) 25, 2078–9 (2009).

R Core Team. R: A Language and Environment for Statistical Computing. R Foundation for Statistical Computing, Vienna, Austria (2015).

Sulakhe, D. et al. Lynx: a knowledge base and an analytical workbench for integrative medicine. Nucleic acids research 44, D882–7 (2016).

Edgar, R., Domrachev, M. & Lash, A. E. Gene Expression Omnibus: NCBI gene expression and hybridization array data repository. Nucleic acids research 30, 207–10 (2002).

